# SugarBase: mapping glycomolecule precursors in microbes

**DOI:** 10.64898/2026.04.20.719630

**Authors:** Jitske M van Ede, Martine C Holst Sørensen, Mark CM van Loosdrecht, Martin Pabst

## Abstract

Glycan biosynthesis relies on nucleotide-activated sugars, essential metabolites across all domains of life, yet their usage in microbes is poorly understood. Here we present SugarBase, a mass spectrometry and bioinformatic pipeline for untargeted exploration of microbial nucleotide sugar networks. SugarBase resolves the chemical complexity of microbial metabolism by combining narrow-window DIA fragmentation with a chemistry-informed parent ion identification algorithm.

Applying SugarBase across a broad phylogenetic range of microbes revealed extensive, species-specific nucleotide sugar profiles, including many candidates with no existing annotation, generating the most comprehensive inventory of nucleotide sugars to date.

SugarBase guided identification of gene clusters and allowed discrimination between pseudaminic- and legionaminic acid–producing strains, where genomic and proteomic data provided only ambiguous information. We resolved distinct nonulosonic acid profiles in several Campylobacter jejuni strains, sugars which may alter susceptibility towards distinct flagellotropic phages. We further identify previously undescribed CMP-activated higher-carbon ulosonic acids in Magnetospirillum, expanding the known chemical space in glycan biosynthesis. In summary, SugarBase supports scalable discovery of microbial nucleotide sugar pathways and enzymes, expanding access to chemically complex glycans and providing new targets for antimicrobial development.

## INTRODUCTION

Nucleotide-activated sugars are central precursors for the diverse range of glycomolecules found across all domains of life^1^. They form storage polymers, structural components, and surface glycans that mediate protection and environmental interactions. In pathogenic bacteria, these glycans often act as virulence factors; for example, O-linked glycans on the flagella of *Campylobacter jejuni* are essential for motility^2-5^. As a result, glycan biosynthesis pathways represent attractive targets for the development of antimicrobials and vaccines^6-9^. Since the discovery of UDP-glucose in the 1950s, many monosaccharides have been identified as UDP, ADP, GDP, CMP, and dTDP conjugates^10^. Their biosynthesis often follows defined biochemical rules, such as the preferential use of specific nucleotides for particular sugars. For example, glucose is predominantly activated in the form of UDP-Glc, which links central metabolism with glycan biosynthesis pathways^11^. While some nucleotide sugars are conserved across taxa, their diversity can vary substantially. Mammalians typically rely on a relatively limited and well-defined pool of nine nucleotide-sugars^1,12^. This diversity expands to more than 30 in plants^13^. In contrast, the prokaryotic repertoire is still largely unexplored and likely includes hundreds to thousands of distinct derivatives.

For example, O-antigen biosynthesis in *E. coli* and *Salmonella* alone involves some 30 different nucleotide sugar intermediates^14^. Furthermore, sugars can be activated by different nucleotides. For example, next to UDP-Glc, also ADP-Glc, GDP-Glc and TDP-Glc are commonly found in microbes, where the different nucleotides likely provide a strategy to partition flux into distinct biosynthetic pathways or classes of glycans^15^.

A few sugar classes are activated exclusively by a single nucleotide. For example, ulosonic acids, including sialic acids and the bacterial analogues pseudaminic and legionaminic acids, have thus far only been observed in CMP-activated form. Sialic acids and their bacterial analogues are important functional sugars involved in recognition and immune evasion and are known virulence factors in pathogenic bacteria. Their exclusive activation is likely dictated by molecular stability, as the α-keto acid group allows a stable linkage to monophosphates, but not to diphosphates^16^.

Currently, only a few complete biosynthetic pathways for nucleotide sugars and their glycans are known in microbes, while the vast majority remain unexplored. Moreover, their chemical synthesis is often infeasible due to stereochemical complexity and limited stability^17,18^, and the lack of commercial standards presents a major bottleneck for developing novel glycoconjugates and biomedical assays. In particular, non-pathogenic microbes represent an attractive source of nucleotide sugars, required to accelerate scalable production of glycoconjugates. For example, CMP-activated nonulosonic acids are particularly challenging to synthesise^1,19,20^, and the biosynthetic routes from *H. pylori* and *C. jejuni* are the most effective ways to produce these compounds^21,22^.

Different analytical strategies are used to identify nucleotide sugars from cell lysates. However, these approaches generally do not support untargeted, large-scale exploration of microbial species. Chromatographic methods, including reverse-phase separations employing ion-pairing reagents and ion-exchange chromatography, are either incompatible with mass spectrometric detection or heavily dependent on the availability of reference standards, limiting their application to complex mixtures of unknown sugars^12,23-26^. By contrast, hydrophilic interaction liquid chromatography and porous graphitic carbon chromatography are compatible with mass spectrometric detection, where the latter is chemically more stable and provides excellent separation of isomeric structures^24,27,28^. Alternatively, capillary electrophoresis was coupled to mass spectrometric precursor ion scanning which allowed an untargeted exploration of nucleotide sugars^29^. However, coupling capillary electrophoresis to mass spectrometry is technically challenging, and high-resolution mass spectrometric analysis of complex cell lysates yields thousands of features, making manual exploration impossible.

Therefore, we developed the SugarBase workflow, which enables untargeted mapping of nucleotide sugar networks in complex, previously uncharacterized microbial species. The workflow combines porous graphitic carbon separation with high-resolution, narrow-window fragmentation. Sugar nucleotides are identified using the SugarBase algorithm, which leverages a large empirical database to link diagnostic fragments to precursor ions, allowing detection of novel derivatives. To demonstrate its performance, we screened a broad taxonomic range of microbes, including mammalian and plant controls, generating the most comprehensive inventory of nucleotide sugars to date.

Interestingly, in several environmental microbes with biotechnological potential, such as *Ca*. Accumulibacter phosphatis and *M. gryphiswaldense*, we identified abundant levels of CMP-activated nonulosonic acids (CMP-NulOs). In addition, we established the CMP-NulO profiles of several pathogenic *C. jejuni* strains, revealing over a dozen different pseudaminic acid (Pse) and legionaminic acid (Leg) derivatives, with the potential to modulate protein and cell surface hydrophobicity, net charge, and susceptibility towards flagellotropic phages. Finally, SugarBase also provides the first evidence of a CMP-activated higher-carbon ulosonic acid discovered in *M. gryphiswaldense*. While differentiation between Leg and Pse is poorly reported in literature, the SugarBase workflow guides identification of biosynthetic gene clusters and allows discrimination between pseudaminic- and legionaminic acid–producing strains, which was validated with orthogonal labelling experiments and synthetic reference standards.

The SugarBase pipeline is freely available as Python code and as a standalone desktop executable with a browser-based graphical user interface that enables interactive exploration of nucleotide sugar profiles: https://sourceforge.net/projects/sugarbase-x/files. All acquired data are publicly accessible through the 4TU.ResearchData platform (https://data.4tu.nl/ project “SugarBase”) and can be reanalysed using the SugarBase workflow.

## MATERIALS AND METHODS

### Sources of microbial biomass and control cells

A table detailing the analyzed biomass, including sources, growth conditions, and suppliers, is provided in SI Excel 1. Specifically, *Campylobacter jejuni, Magnetospirillum gryphiswaldense*, and *Ca*. Accumulibacter phosphatis were cultured as described in the following. **Culturing of Campylobacter jejuni**. *C. jejuni* strains NCTC12658 and NCTC12662 were grown in liquid Brain Heart Infusion broth under microaerophilic conditions (6% CO2, 6% O2, 88% N2) at 41.6°C. Biomass was harvested at an OD_600_ of 0.25 by centrifuging at 6500 rpm, for 7 min, in multiple rounds. Campylobacter *jejuni* strains 11168 (DSM 27585) and 81116 (DSM 24189) were cultured on Mueller–Hinton agar supplemented with Oxoid™ Campylobacter Growth Supplement (Thermo Fisher Scientific). Microaerophilic conditions were generated in a 2.5 L anaerobic jar using Anaerocult™ C (Merck). Plates were incubated for 48 h at 37°C. Biomass was harvested with sterile cotton swabs into sterile 1× PBS, centrifuged (10 min, 14,000 rpm, 4°C), and stored at −80°C until use. **Culturing of *M. gryphiswaldense***. *Magnetospirillum gryphiswaldense* DSM 6361 was cultured in a lactate-containing medium under microaerophilic conditions at 30 °C. 5 L culture in 20 L bottle was initiated under the gas phase containing 2% O_2_ in argon to support growth onset. Upon visible initiation of growth, microaerophilic conditions were relaxed by opening the culture flasks to atmospheric air through a sterile filter and the bottle was placed on a rotary shaker at 100 rpm, allowing continued growth under oxic conditions. ***Ca*. Accumulibacter enrichment**. A laboratory-scale enrichment of *Candidatus Accumulibacter* was maintained in sequencing batch reactor mode under alternating anoxic/aerobic feast–famine cycles following methods described recently^30^. Briefly, the enrichment was maintained at 20°C and pH 7.0 and fed with acetate as the primary carbon source. Anoxic phases were maintained by N_2_ sparging, followed by aerobic phases with air supply. Biomass was harvested at the end of the aerobic phase. **Metabolite extraction from cell materials**. 500 µL pre-cooled extraction solvent solvent (-20°C, 40% acetonitrile, 40% methanol, 20% MS-grade water) and 0.1 g acid-washed glass beads (500 µm, Sigma Aldrich) were added to 100–200 mg of biomass, wet weight. Cells were disrupted by three cycles of vortexing for 1.5 min, with 1 min on ice between cycles. Plant tissues were ground in liquid nitrogen using a mortar and pestle prior to solvent addition. The lysates were centrifuged at 14000 rpm, 4°C for 15 min in an Eppendorf container (5424R), and supernatants were collected. The extracts were purified and concentrated using HyperSep™ Hypercarb™ SPE 96-well plates (Thermo Scientific, Germany). In short, the columns were conditioned with 600 µL 10 mM ammonium bicarbonate (ABC) in 60% acetonitrile (ACN) and equilibrated with 2 × 600 µL MS-grade H_2_O. Samples were diluted 1:10 with MS-grade H_2_O, loaded onto the columns, washed with 1 mL MS-grade H_2_O, and eluted with 2 × 200 µL 10 mM ABC in 60% ACN. Eluates were divided into three equal aliquots and dried in a SpeedVac concentrator at room temperature. Dried extracts were stored at −80°C and reconstituted in 15 µL MS-grade H_2_O prior to mass spectrometric analysis.

### SugarBase workflow. A. PGC chromatography coupled to narrow-mass window DIA

Data independent mass spectrometeric analysis was performed using an Acquity Ultra Performance Liquid Chromatograph (Waters, UK) coupled to a Q Exactive™ Focus Hybrid Quadrupole-Orbitrap™ Mass Spectrometer (Thermo Scientific, Germany). Chromatographic separation was performed on a 100 × 1 mm Hypercarb™ Porous Graphitic Carbon Column (Thermo Scientific, No. 35005-101030) at a flow rate of 100 µL/min. Additional electrical grounding between the separation column and the electrospray ionization source was applied to prevent interference of the electrospray voltage with chromatographic separation^31^. Mobile phase A consisted of 15 mM ammonium bicarbonate in MS-grade H_2_O and mobile phase B of 100% acetonitrile. A linear gradient from 2.5% B to 35% B was applied over 18 min, followed by a linear gradient to 60% B over another 6 minutes. Each sample was analysed in duplicate followed by two blanks. The mass spectrometer was operated in parallel reaction monitoring mode, where a mass range of 500–612 Da (Part 1) was fragmented continuously in 8 Da steps, using an isolation window of 10 m/z. Each window was measured twice, first applying a NCE of 10 (pseudo MS1) followed by a NCE of 28 (MS2). Electrospray ionization was performed in negative ionisation mode. MS analysis was performed at a resolution of 35 K, with a AGC target of 2.00E5 and a maximum injection time of 75 ms. Data were acquired in the centroid mode, with microscans set to 1. Every sample was injected a second time using an extended mass range, covering 596–708 m/z (Part 2). Raw data were analysed using XCalibur 4.1 (Thermo Fisher Scientific, Germany), Matlab R2023b (MathWorks) and Python 3.9. The mass spectrometer was calibrated using the Pierce™ LTQ ESI negative ion calibration solution (Thermo Fisher Scientific, Germany). **B. Nucleotide sugar data processing pipeline**. Potential nucleotide-sugars were identified using the Python SugarBase pipeline. Briefly, raw mass spectrometry files were converted to “.mzXML” format using msConvert^32^. Peak lists were imported from which mass lists were extracted. The odd and even scans were separated, representing pseudo-MS1 and fragmented MS2 data, hereafter referred to as “MS1” and “MS2,” respectively. “MS1” data was filtered to exclude masses outside the designated isolation window (“channel”) to remove potential in-source fragments. Peaks with an intensity <1000 or peaks outside the empirical retention time window (4.5–15.0 min) were not further processed. Doubly charged peaks were removed, and deisotoping was performed considering 10 ppm mass error. Up to three isotopes peaks with an abundance ratio threshold compared to M were considered (M+1<0.75, M+2<0.33, M+3<0.15). A cell array was generated for each “MS2” channel (10 Da window), containing scan index, mass channel, retention time, fragment peaks, and peak intensities. MS2 arrays containing potential nucleotide fragment markers (CMP, UMP, UDP, GMP, GDP, AMP, ADP, CDP, dTMP, dTDP, TMP, and TDP; masses 322.0446, 323.0286, 402.9949, 362.0507, 442.0171, 346.0558, 426.0221, 402.0109, 321.0493, 401.0157, 337.0442, and 417.0106, respectively) were extracted considering a mass error of 10 ppm. Corresponding MS1 scans were then examined for candidate nucleotide sugar peaks containing the identified nucleotide marker. Discrimination between true nucleotide sugars and background metabolites was achieved using a sugar composition database based on work published by Pabst and co-workers (2021)^33^. All possible combinations of C_5-16_H_8-27_N_0-4_O_3-9_S_0-1_ were generated and constrained using a series of physicochemical filters. Briefly, compositions with monoisotopic masses below 100 Da or above 400 Da were excluded. Mass defect thresholds were set to exclude values below 0.01 or above 0.19, with additional constraints applied for lower mass species. Compositions with masses between 100–200 Da and mass defects greater than 0.09, and those between 100–250 Da with mass defects greater than 0.13, were excluded. Double bond equivalents (DBE) were restricted to values between 1.0 and 6.0, with compositions in the 100–150 Da mass range excluded if DBE exceeded 3.0. Elemental ratio filters were applied by excluding compositions with carbon-to-hydrogen (C/H) ratios below 0.40 or above 1.25, and carbon-to-(oxygen plus nitrogen) [C/(O+N)] ratios below 0.90 or above 2.10, accounting for potential sulfonic acid formation. Finally, nitrogen and sulfur content was limited by excluding compositions with more than one nitrogen atom in the 100–200 Da mass range, more than three nitrogen atoms in the 100–275 Da range, or any compositions containing both nitrogen and sulfur atoms. The resulting sugar compositions were then combined with the nucleotide masses of CMP, UDP, GDP, ADP, CDP, dTDP, and TDP, to generate a theoretical nucleotide sugar database. All MS1 mass peaks matching the corresponding nucleotide sugar space within 10 ppm mass window were retained for further processing. Precursors adjacent to CMP marker fragments were matched to CMP and CDP sugars. Moreover, to increase robustness, only precursor ions detected in at least three consecutive scans within the same channel and exhibiting the corresponding nucleotide marker fragment in MS2 spectra were retained for further processing. Finally, to provide confidence in the nucleotide sugar candidate outputs, candidates were scored using multiple criteria. Briefly, a rule-based scoring system was employed, where hits detected in at least four consecutive scans were assigned a score of +10. Signal consistency across scans was further evaluated using Pearson correlation coefficients calculated over three consecutive scans. The “MS1” scan with the highest intensity for a given nucleotide-sugar was designated the “main hit” provided that both the precursor and fragment masses were present in the adjacent spectra of the same channel. These three spectra were used to calculate the Pearson correlation coefficients, where correlations greater than 0.80 were rewarded (+10), whereas correlations below 0.70 and 0.50 were penalized (−10 and −50, respectively). For non-CMP-activated nucleotide sugars, the ratio of precursor ion intensity in the MS1 scan to fragment ion intensity in the corresponding MS2 scan was used as an additional criterion. Ratios below 100 were rewarded (+10), ratios below 1 were penalized (-10), and ratios exceeding 110 were strongly penalized (-50). For CMP-activated nucleotide sugars, scoring criteria included the ratio of precursor intensity in the MS2 scan relative to the CMP fragment intensity in the same MS2 scan, with values ≤25 rewarded (+10) and values >25 penalized (−50). High precursor abundance in MS1 (>100K intensity units) and presence of a characteristic CO_2_ neutral loss in MS2 spectra (-CO_2_, -43.98983) contributed positively (+10). In addition, an elemental parity (“N-rule”) was applied, requiring both nitrogen and hydrogen atom counts to be either even or odd. Deviations from this rule resulted in a penalty (−50). Finally, for each nucleotide sugar candidate, an extracted ion chromatogram was generated together with the corresponding precursor and fragmentation spectra. These results are visualized via the browser-based interface and exported as PNG files, while a summary table containing scores and assigned chemical compositions is reported as an Excel file. Additional targeted nucleotide sugar analyses from acquired data were performed by specifically screening for common pentose, hexose, and heptose monosaccharide derivatives, including amino sugars and N-acetylated, deoxy, dideoxy, uronic acid, and N-acetyluronic acid variants. These monosaccharides were combined *in silico* with the common activating nucleotides ADP, UDP, GDP, dTDP, TDP, CDP, and CMP. In addition, CMP-, UMP-, GMP-, dTMP-, and TMP-activated NeuAc, NulOAc_2_, NulOAcAm, and NulOAcAla, were included (SI Excel file, targeted analysis). The same filtering and scoring criteria as used for untargeted discovery were applied, except that no MS1 intensity threshold was used and only candidates with scores ≥0 were retained for further analysis. **C. SugarBase code availability**. The SugarBase Python code and the standalone desktop executable with graphical user interface are provided via SourceForge: https://sourceforge.net/projects/sugarbase-x/. The provided pipeline was established and tested using Q Exactive Orbitrap data (Thermo Fisher Scientific).

### Shotgun proteomic analysis

Approximately 25 mg (wet weight) of each cell biomass material was resuspended in 175 µL 50 mM TEAB containing 1% (w/w) NaDOC, adjusted to pH 8 and 175 µL Bacterial Protein Extraction Reagent (B-PER; Thermo Scientific, cat. 78243). After addition of 0.15 g acid-washed glass beads (150–212 µm; Sigma-Aldrich), samples underwent three cycles of bead beating (1.5 min) using a bench top vortex, with 1 min breaks on ice in between. Subsequently, the samples were centrifuged at 14000 rcf for 10 min at 4 °C. For *C. jejuni*, the samples were additionally incubated at 80°C for 3 min at 1000 rpm, centrifuged, and supernatants were passed through a 0.22 µm filter before further processing. For all samples, proteins were precipitated from the supernatants by adding TCA at a 1:4 ratio, vortexed, and incubating at 4°C, followed by centrifugation to collect the protein pellets. Pellets were washed with ice-cold acetone and resuspended in 200 µL 6 M urea (*C. jejuni*) or 100 µL 6 M urea (*E. coli* and *Ca*. Accumulibacter) and incubated at 37°C, 1000 rpm until solubilized. Reduction was performed by adding 10 mM dithiothreitol in 200 mM ammonium bicarbonate to a final DTT concentration of 2.3 mM and incubating for 1 h at 37°C with shaking at 300 rpm. Alkylation was carried out by adding 20 mM iodoacetamide in 200 mM ammonium bicarbonate to a final concentration of 3.75 mM and incubating for 30 min at room temperature in the dark. Samples were diluted with ammonium bicarbonate buffer to bring the sample to below <1 M urea. A 100 µL aliquot for each sample was digested with 5 µL of 0.1 µg/µL trypsin (Promega, cat. V5111) overnight at 37°C, 300 rpm. On the following day, samples were cleaned using Oasis HLB 96-well µElution plates (2 mg sorbent, 30 µm, Waters, cat. 186001828BA). Columns were conditioned with 750 µL methanol, equilibrated with 2 × 500 µL MS-grade water, and loaded with samples. After washing with 350 µL 5% methanol in MS-grade water, peptides were eluted with 200 µL 2% formic acid in 80% methanol followed by 200 µL 1 mM ABC in 80% methanol. Eluates were dried in a SpeedVac concentrator and reconstituted in 3% acetonitrile containing 0.1% formic acid to a final concentration of approximately 500 ng/µL. 1 µL of proteolytic digest, as estimated by NanoDrop UV spectrophotometer at 280 nm, was used for shotgun proteomic analysis, using an EASY nano-LC 1200 system coupled to a Q Exactive Plus Orbitrap mass spectrometer (Thermo Scientific, Germany). Peptides were separated on a 0.05 × 150 mm C18 analytical column (Thermo Scientific, catalog no. 164943). Mobile phase A consisted of 0.1% formic acid and 1% acetonitrile in MS-grade water, while mobile phase B contained 0.1% formic acid and 80% acetonitrile in MS-grade water. The LC gradient started with a 5 min hold at 5% mobile phase B, followed by a linear increase to 30% B over 90 min, then ramped to 75% B over an additional 25 min, maintaining a constant flow rate of 300 nL/min throughout the run. Full MS scans were acquired at a resolution of 70K with an automatic gain control target of 3E6 and a maximum injection time of 100 ms. Data-dependent acquisition was employed, where top 10 precursor ions were selected across a mass range of 550–1500 m/z using an isolation window of 1.7 m/z. Fragmentation was performed using a normalized collision energy of 28%. Fragmentation spectra were recorded at a resolution of 17.5K with an AGC target of 5E5 and a max injection time of 100 ms. Protein identification was carried out using PEAKS Studio X (Bioinformatics Solutions Inc., Canada). Database searches were performed against protein sequence databases obtained from UniProt for Campylobacter jejuni (UP000000799) and Escherichia coli K-12 (UP000000625), together with the GPM cRAP contaminant database. The reference proteome for *Ca. Accumulibacter phosphatis* was derived from whole-metagenome sequencing, as described previously^34^. Carbamidomethylation was specified as a fixed modification, whereas oxidation and deamidation were included as variable modifications. Trypsin was selected as the digestion enzyme, permitting up to three missed cleavage sites. Mass tolerance settings were defined as 20 ppm for precursors and 0.02 Da for fragment ions. Peptide and protein identifications were filtered at a false discovery rate (FDR) of 1%, and protein identifications were considered significant when supported by at least two unique peptides. The NovoBridge and NovoLign pipelines were used to confirm the taxonomic purity of the samples^34,35^. **DMB labelling and mass spectrometric analysis of free NulOs**. Approximately 50 mg (wet weight) of biomass was suspended in 250 µL of 2 M acetic acid, followed by the addition of 0.15 g acid-washed glass beads (150–212 µm). Cells were disrupted by three bead-beating cycles (1.5 min each), with 1 min incubation on ice and 2 min centrifugation (14,000 rpm, 4°C) between cycles. Acid hydrolysis was performed at 80°C for 2 h at 300 rpm using an Eppendorf heating block. After centrifugation (10 min, 14,000 rpm, 4°C), supernatants were filtered through a 0.22 µm membrane to remove residual cells and dried in a vacuum concentrator at 60°C. Alternatively, dried nucleotide-sugar extracts, obtained as described in the *Metabolite extraction* section, were hydrolyzed in 200 µL 2 M acetic acid under identical conditions and dried at 60°C. For DMB derivatization, dried hydrolysates were resuspended in DMB labeling solution (1.4 M acetic acid, 0.75 M 2-mercaptoethanol, 18 mM sodium dithionite, 7 mM DMB) and incubated for 2.5 h at 50°C (600 rpm), using 50 µL for biomass-derived samples and 25 µL for nucleotide-sugar extracts. DMB-labeled samples were analyzed using the protocol described in Pabst et al.^33^ Briefly, labeled lysates were separated on a Waters Acquity UPLC system equipped with a C18 BEH column (1.7 µm particle size) and coupled to a Q Exactive Focus Orbitrap mass spectrometer. The instrument was operated in positive electrospray ionization mode with alternating full MS scans and narrow-mass window fragmentation scans covering the m/z range 390–525. Mass windows of 5.5 Da were sequentially isolated and fragmented using a normalized collision energy of 26. Raw mass spectrometric data were processed using MATLAB scripts as described by Kleikamp *et al*.^36^ Additional MS3 fragmentation experiments were performed on DMB-labeled ulosonic acids extracted from *Magnetospirillum gryphiswaldense* and on DMB-labeled synthetic standards of legionaminic acid and Neu5Ac. Compounds were separated using an UltiMate 3000 LC system equipped with an HT µPAC Neo Plus C18 column (Thermo Fisher Scientific) and coupled to an Orbitrap Eclipse Tribrid mass spectrometer, operated in positive ionsiation mode. The flow rate was set to 1.5 µL/min, with water plus 0.1% formic acid as solvent A, and 80% acetonitrile, 0.1% formic acid as solvent B. A gradient from 7.5% to 45% solvent B was maintained over 15 min. The mass spectrometer continuously targeted the intact DMB-XulO, DMB-LegAc_2_, and DMB-Neu5Ac precursor ions, followed by targeted fragmentation of the corresponding C9 and C10 marker fragments m/z 295.0716, 297.0872, and 311.1056, respectively. MS2 isolation was performed in the quadrupole with a 1.5 m/z window using a normalized collision energy of 25 and Orbitrap detection at 500K resolution. MS3 isolation was carried out in the linear ion trap using a 2 m/z window and a normalized collision energy of 32 or 40 and Orbitrap detection at 240K resolution. Fragmentation spectra were manually interpreted using Xcalibur, ChemDraw, and ChemCalc^37^. **Sequence alignment, and investigation of biosynthetic gene clusters (BGCs)**. Reference protein sequences representing the legionaminic acid (Leg) and pseudaminic acid (Pse) biosynthetic pathways were compiled to identify homologous genes across other microbial genomes. For the legionaminic acid pathway, reference proteins included LegB (Q0P8T8), LegC (Q0P8T7), LegH (Q0P8V9), LegG (Q0P8T0), LegI (Q0P8T1), LegF (Q0P8S7), Maf4/1 (Q0P8S3), PtmG (Q0P8T3), PtmH (Q0P8T2), and GlmU (Q9PPA2). UDP-GlcNAc–linked pathway components included PtmE (Q0P8S9), PgmL (Q0P8K7), PtmA (Q0P8S6), and PtmF (Q0P8S8). For the pseudaminic acid pathway, reference sequences comprised PseB (Q0P8W4), PseC (Q0P8W3), PseH (Q0P8U4), PseG (Q0P8U5), PseI (Q0P8U0), PseF (Q0P8U6), and PseE (Q0P8S2). Additional bacterial nonulosonic acid biosynthesis references included PglF (Q0P9D4), PglE (UniRef100_Q0P9D3), PglD (UniRef100_Q0P9D1), NeuC/Lpg0753 (Q5ZXH8), NeuB (Q5ZXH9), and NeuA (A0A0P0LXX4). These reference sequences were used to construct a custom database for homology searches using DIAMOND. Predicted proteomes from the analyzed microbial genomes were aligned against this reference set using DIAMOND with default parameters to identify candidate pseudaminic acid and legionaminic acid–associated biosynthetic genes and homologous pathway components. Alignments within a 5% range of the top alignment score for a query were reported (‘--top 5’). DIAMOND alignment results were processed using a custom Python script to retain high confidence homologs. For each alignment, query coverage was approximated from the aligned query span, and hits were filtered using stringent criteria to exclude spurious or local matches. Only alignments covering at least 70% of the query sequence were retained to avoid partial or domain-level hits. In addition, a minimum sequence identity of 25% was required, together with a minimum bitscore of 80 and an E-value threshold of ≤1×10^-10^ to ensure high statistical significance. Analysis of biosynthetic gene clusters was performed using the NCBI Genome Viewer. For *Ca*. Kuenenia stuttgartiensis, analyses were based on the NCBI reference sequence NZ_LT934425.1, and for *Campylobacter jejuni* subsp. *jejuni* NCTC 11168 (ATCC 700819), on the reference sequence NC_002163.1. **Synthetic sugar standards and 13C labelled metabolite extract**. 13C-labeled *Penicillium chrysogenum* extract was produced in-house as described previously^38^. UDP-Glc was obtained from Sigma-Aldrich (Cat. No. U4625-100MG), UDP-GlcNAc from Sigma-Aldrich (Cat. No. U4375-100MG), CMP-Neu5Ac from BioSynth (Cat. No. MC04391), Pse5,7Ac_2_ from BioSynth (Cat. No. MP07650), and Leg5,7Ac_2_ from Synvenio (Cat. No. SV7243). The AdvanceBio Sialic Acid Reference Panel was obtained from Agilent (Cat. No. GKRP-2503), and KDN from Sigma-Aldrich (Cat. No. 60714).

## RESULTS AND DISCUSSION

### SugarBase: untargeted exploration of nucleotide sugar networks in microbes

The complexity of crude cell lysates, the chemical diversity of microbial nucleotide sugars, and the high data density of next-generation mass spectrometers require open searches and advanced data mining strategies. To address this, we developed a mass spectrometry workflow that combines porous graphitic carbon (PGC) separation, continuous narrow-window DIA scanning, and a nucleotide sugar discovery software (Figure 1). PGC allows analysis of crude lysates while providing excellent isomer separation of polar compounds. Narrow mass windows (over an m/z range of 500–708, negative mode) are sequentially scanned to maximize sensitivity for phosphate-containing nucleotide sugars. Each window is acquired twice: first at low collision energy to generate a focused pseudo-MS1 spectrum of precursor ions, and then at higher collision energy to obtain corresponding fragmentation spectra (“true MS2”). Narrow isolation windows in fast mass spectrometers reduce spectral complexity by limiting the number of co-fragmented features. In microbial systems, the nucleotide moiety is known which can be either CMP, UDP, GDP, ADP, CDP, dTDP or TDP, whereas the attached sugar is often unknown. To identify such precursors, we developed an algorithm that links nucleotide marker fragments to the correct MS1 precursor ions by matching against a large theoretical sugar composition database containing >2200 entries, allowing assignment of the sugar and nucleotide components (Figure 1, SI Excel 2, SI DOC). To avoid false positives, we implemented a range of inclusion and scoring criteria as detailed in the SI DOC. For example, nucleotide sugars and their marker fragments were expected to be present in at least three consecutive scans within the same isolation window, and co-elution of precursor and marker fragment was confirmed using Pearson correlation. Additionally, precursor intensity, and fragment-to-precursor abundance ratios were expected to fall within empirical ranges. For CMP-nonulosonic acids, additional scoring criteria were included, including the presence of CO_2_ loss (from the carboxyl group) and consistency with the nitrogen rule, where the nominal mass reflects the expected number of nitrogen atoms.

**Figure 1.**
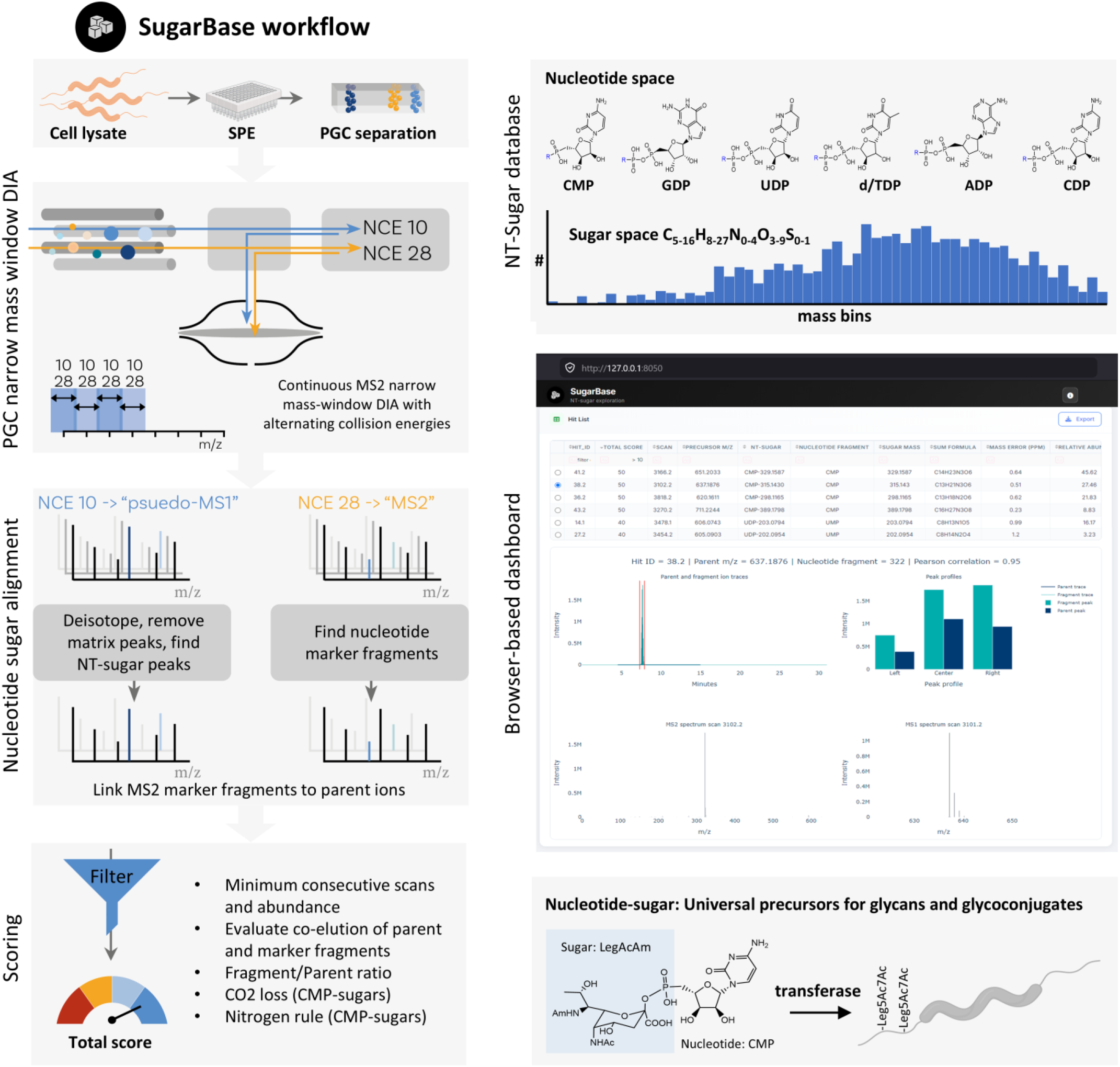
SugarBase: mass spec and bioinformatics workflow for untargeted exploration of microbial nucleotide sugars networks. a) Metabolite extracts are analyzed by porous graphitic carbon separation coupled to narrow-window DIA mass spectrometry. Each window is first acquired at low collision energy to record intact precursor ions (pseudo-MS1), followed by higher energy to obtain fragment spectra (MS2). Pseudo-MS1 spectra are deisotoped and filtered, while MS2 spectra are screened for nucleotide marker fragments. Matching scans are linked to identify nucleotide sugars, and precursor ions are assigned using a sugar composition database. Candidates are evaluated based on consecutive detection, co-elution of precursor and marker ions, realistic intensity ratios, and for CMP-linked sugars, CO_2_ neutral loss and the nitrogen rule. Identifications can be explored via a browser-based Dash interface, of which a screenshot as illustration is displayed in the figure. Structures of CMP, CDP, UDP, dTDP, TDP, ADP, and GDP are shown (reported as [M–H]^-^ ions). The histogram below shows the theoretical composition space of nucleotide-linked sugars, generated from all plausible C_5-16_H_8-27_N_0-4_O_3-9_S_0-1_ combinations, yielding >2200 candidates. These are used to annotate precursor ions when a nucleotide marker is detected. Finally, an example of a nucleotide-sugar as precursor for glycoconjugate is illustrated, showing the transfer of Legionaminic acid onto the flagella of *Campylobacter jejuni*.

The SugarBase tool visualizes identified nucleotide sugars in a browser-based dashboard, which enables interactive exploration of candidate features, scores, mass traces, and extracted ion chromatograms. The SugarBase open-source Python code, standalone desktop application, and documentation are freely available via SourceForge. While the current pipeline was developed and tested using Q Exactive Orbitrap (Thermo Fisher Scientific) data, all parameters are adjustable to support other acquisition methods and vendors, provided raw files can be converted to mzXML format and meet the required input structure.

### Highly divergent nucleotide sugar profiles across microbes

To demonstrate the SugarBase workflow we mapped the sugar profiles across a broad phylogenetic spectrum of microbes, encompassing environmental, pathogenic, and biotechnologically relevant taxa (SI Excel 1), including *Campylobacter, Pseudomonas, Escherichia, Streptomyces, Magnetospirillum, Legionella*, anammox bacteria, archaea, and multiple enrichment cultures. We furthermore included well-studied eukaryotic samples as controls, for example, mammalian cells were included as these produce a defined set of nucleotide sugars, and plant species were included as these are devoid of CMP-NulOs.

Interestingly, for prokaryotes, the principles that govern sugar-type usage and nucleotide activation preferences remain largely unknown^39^. Therefore, we first systematically screened these microbes for the presence of a systematic matrix of nucleotide/sugar combinations.

We generated this matrix of nucleotide sugars by combining common sugar chain lengths with typical modifications (oxidation, deoxygenation, and amination) across all major nucleotides (UDP, ADP, GDP, CDP, CMP, dTDP, and TDP). Nonulosonic acids were paired exclusively with monophosphate nucleotides (UMP, AMP, GMP, CMP, dTMP, and TMP), as these form only stable bonds with monophosphates^16^. This produced a matrix of 203 nucleotide sugar combinations (SI Excel 2), which was used for targeted searches with the SugarBase pipeline, applying default filtering and scoring parameters without abundance cutoffs.

This provided a view on nucleotide sugar usage and combination across a broad phylogenetic spectrum. Of the 203 theoretical nucleotide sugars, 33 were detected in more than one sample (>50 when including single detections, Figure 2, SI Excel 3 & 4, SI DOC). The mammalian and yeast controls showed the expected set of sugars and activating nucleotides, such as abundant UDP-hexose, UDP-HexNAc, GDP-hexose, GDP-deoxyhexose, dTDP-hexose, ADP-pentose, ADP-deoxyhexose, and CMP-Neu5Ac in mammalian cells (right columns, Figure 2a). The abundant ADP-pentose peak likely represents ADP-ribose derived from nicotinamide cofactors, that is involved in ADP-ribosylation and DNA repair rather than glycomolecule biosynthesis.

**Figure 2.**
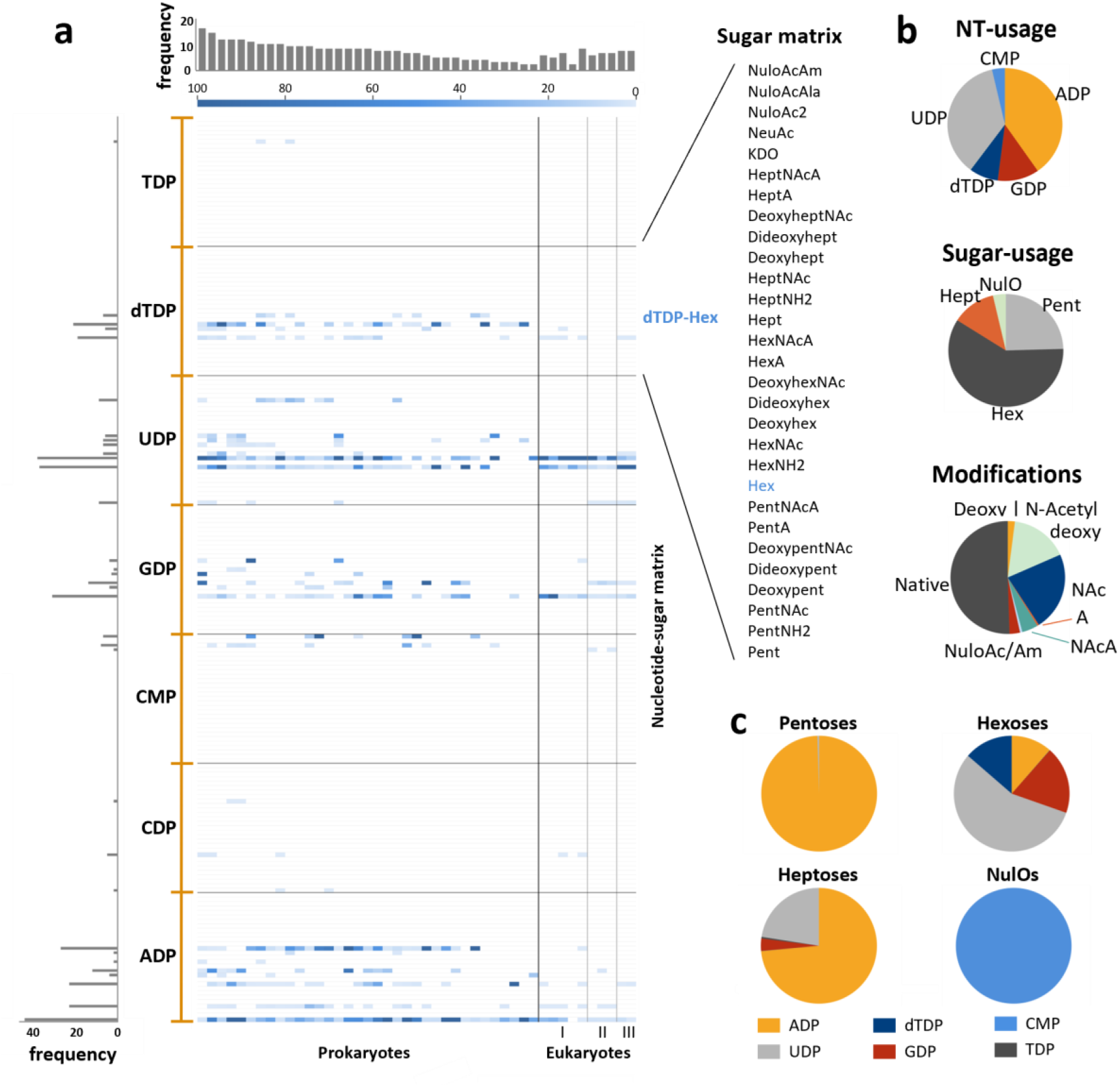
Systematic survey for nucleotide sugar combinations across a broad phylogenetic spectrum. a) The heatmap shows the outcome of a targeted search using a systematic matrix of nucleotide sugar combinations across a broad phylogenetic spectrum of microbes, with mammalian and yeast samples included as controls. A matrix of 203 nucleotide sugars combined common pentose, hexose, heptose, and nonulosonic acid sugars and their modifications to the seven activating nucleotides UDP, ADP, GDP, CDP, CMP, dTDP, and TDP. The monosaccharide classes of the detected nucleotide sugar candidates are displayed on the y-axis, and the color scale represents normalized peak intensities. The bar graph on the left of the heatmap shows the detection frequency of individual nucleotide sugars across all microbes, while the bar graph on the top shows the number of identified nucleotide-sugars per organism. I: yeast; II: mammals; III: plants. B) The pie charts show the nucleotide activation preference, the sugar class usage and the modification usage across all analyzed samples. c) The pie charts show nucleotide activation preferences for each sugar type (pentoses, hexoses, heptoses, and nonulosonic acids) across all analyzed samples.

In contrast, prokaryotes displayed a diverse spectrum of nucleotide sugars with many species-specific features^36^. Generally, ADP- and UDP-linked sugars dominated, dTDP-linked sugars were also frequently observed, but CDP activation was only observed in a few cases (Figure 2). Pentoses were predominantly ADP-linked, while hexoses showed greater heterogeneity and occurred as UDP-, ADP-, GDP-, and dTDP-linked sugars (Figure 2c). The different nucleotides may partition the carbon flux into different biosynthetic glycan pathways, as glycosyltransferases are selective for the linked nucleotide^15,40^. Nonulosonic acids were found exclusively linked to CMP, with no significant evidence for linkage to other nucleotides (Figure 2c). While UDP-Hex, UDP-HexNAc, and GDP-Hex are expected to be universal metabolic precursors, in prokaryotes they were not always the most abundant and in some cases were not detected at all. Compared to eukaryotes, this variability likely reflects the greater metabolic flexibility of microbes and the diversity of carbon sources across their environments.

### Deep untargeted mapping of nucleotide sugars in pathogens and microbial model organisms

Next, we employed SugarBase to perform a fully untargeted profiling to also capture previously uncharacterized nucleotide sugars and to explore CMP-NulO modification profiles in common pathogens. Thereby we identified 87 recurring nucleotide sugars (using a score cutoff of ≥20 and detection in at least two species, Figure 3a, SI Excel 5). Notably, 10 of the 25 most frequent features could not be assigned to known nucleotide sugars. However, we also observed many recurring background features, including AMP and AMP-pentose related fragments, likely originating from in-source fragmentation or degradation of NAD- and CoA-derived metabolites. All commonly observed background feature are summarized in SI Excel 3. Overall, the most frequent nucleotide sugars (detected in >5 organisms) included central precursors for glycan and cell envelope biosynthesis, such as UDP-HexNAc, UDP-Hex, GDP-Hex, UDP-MurNAc, ADP-heptose, ADP-deoxypentose, ADP-hexose, dTDP-deoxyhexose, GDP-deoxyhexose, dTDP-hexose, UDP-pentose, and various CMP-NulOs. Notably, several less studied but abundant species including CDP-Hex, UDP-deoxyHexNAc, UDP-Hexdi/triNAc, and HexNAc derivatives linked to dTDP, ADP, or GDP, were also observed, which shows the metabolic flexibility in microbes. Furthermore, Figure 3b zooms in on microbes with biotechnological relevance. The model unicellular eukaryote and cell factory *S. cerevisiae*, showed mostly UDP-HexNAc, followed by UDP-Hex, GDP-Hex, ADP-ribose, and some lower abundant dTDP-hexose derivatives, in line with known fungal metabolic routes (Figure 3b). Similarly, the embryonic kidney (HEK) cells, a well-characterized mammalian model, showed nucleotide sugars consistent with the conserved eukaryotic glycosylation pathways (SI Excel 5)^12,41^. In contrast, bacterial samples showed much greater heterogeneity, both within and between species. The pathogen *Campylobacter jejuni* produces abundant GDP-heptose and multiple CMP-linked sugar derivatives alongside UDP-HexNAc and UDP-dHexNAc, consistent with bacillosamine-containing N-glycan and lipooligosaccharide biosynthesis pathways^42,43^. *Ca*. Kuenenia stuttgartiensis, an anammox bacterium involved in global nitrogen cycling and wastewater treatment^33^ displayed multiple CMP-linked sugars, UDP-glucuronic acid, UDP-HexNAcA, UDP-pentose, ADP- and GDP-linked heptoses, and UDP-MurNAc. Interestingly, Ca. Kuenenia stuttgartiensis activates heptose with both ADP and GDP, where the ADP-heptose is used in lipopolysaccharide biosynthesis^44^, and GDP-heptose likely relates to the recently reported oligoheptosidic glycans modifying the surface layer^33^. The magnetotactic bacterium *Magnetospirillum gryphiswaldense*, a model platform in nanobiotechnology^45^, showed one of the most distinctive nucleotide sugar profiles, with several previously uncharacterized CMP-linked sugars further described in the following section. A particularly complex spectrum was observed in the *Ca*. Accumulibacter phosphatis enrichment, which next to its mixed composition is also known for its metabolic versatility^46^, producing unique compounds such as UDP-HexNAc2A, ADP-dHex, and dTDP-dHex along with UDP-HexNAcA and ADP-hexose. In many species we also detected abundant UDP-MurNAc and downstream UDP-muramyl peptide intermediates from peptidoglycan and cell wall assembly^47^. Interestingly, the analysed microbes frequently showed CMP-, GMP-, or ADP-linked sugars that were more abundant than the otherwise central UDP-Hex and UDP-HexNAc derivatives. Several nucleotide sugars with high confidence scores could not be annotated, highlighting the unexplored diversity of microbial metabolism.

**Figure 3.**
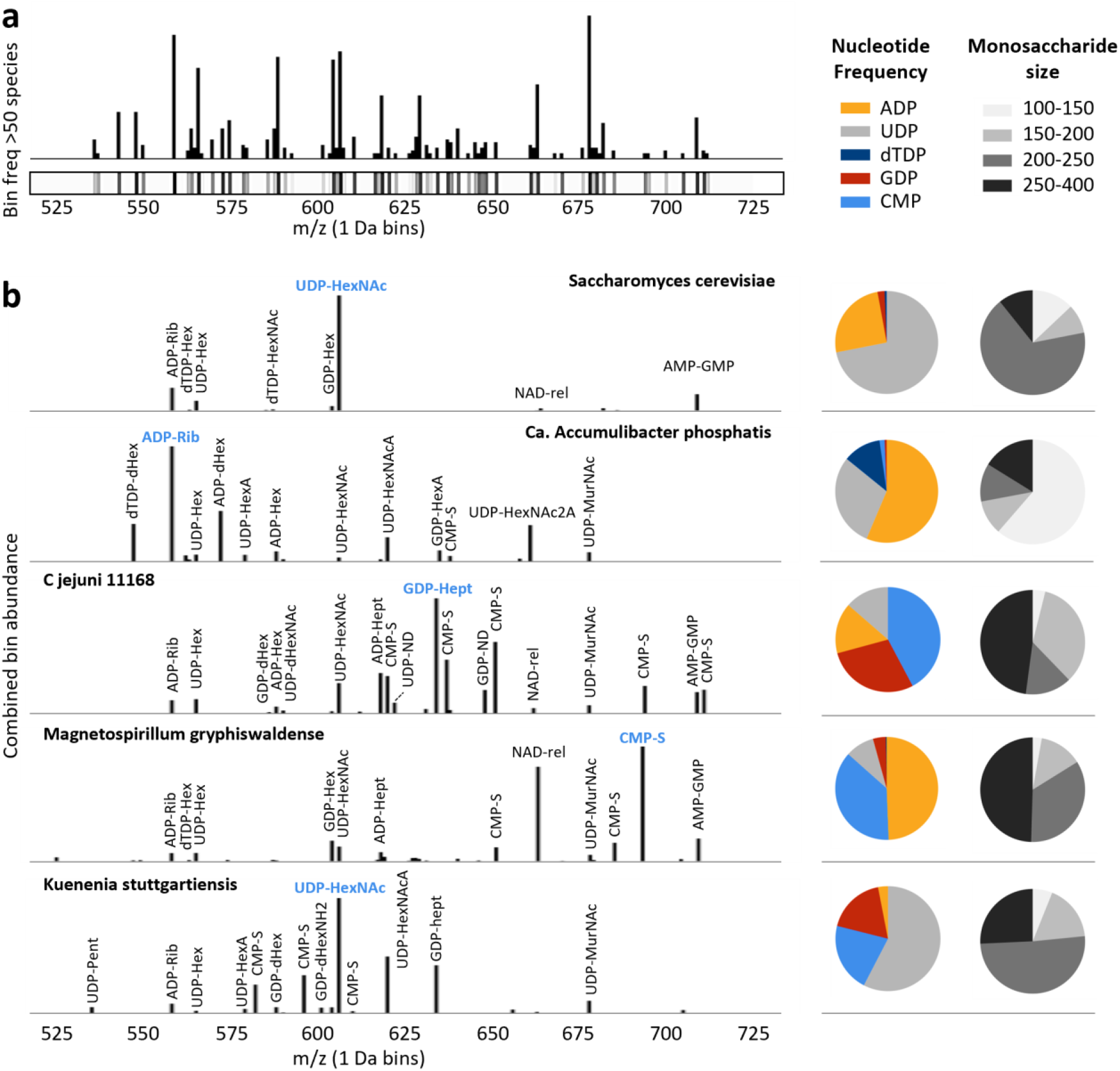
Fully untargeted SugarBase workflow shows diverse nucleotide sugar profiles across pathogens and biotech strains. a) The frequency histogram and heatmap show all nucleotide sugars identified across the analyzed microbes using the fully untargeted SugarBase workflow. b) The histograms below show nucleotide sugar profiles for individual species, including *Saccharomyces cerevisiae, Ca*. Accumulibacter phosphatis, *Campylobacter jejuni* NCTC 11168, *Magnetospirillum gryphiswaldense*, and *Ca*. Kuenenia stuttgartiensis. The pie charts next to each spectrum indicate the relative abundance of nucleotide types: ADP, UDP, CDP, (d)TDP, GDP, CMP and monosaccharide size classes: 100–150, 150–200, 200–250, and 250–400 Da. A summary table of all identified nucleotide sugar candidates across species, along with detailed tables for each organism, is provided in SI Excel Table 5. Abbreviations: CMP-S, CMP-sugar; dHex, deoxyhexose; dHexNAc, N-acetyl-deoxyhexose; Hept, heptose; Hex, hexose; HexA, hexuronic acid; HexNAc, N-acetyl-hexose; HexNAcA, N-acetyl-hexuronic acid; HexNAc2A, di-N-acetyl-hexuronic acid; MurNAc, N-acetyl-muramic acid; NAD-rel, NAD-related compound; ND, not defined; Pent, pentose; Rib, ribose.

### Mapping CMP-NulO modification profiles in pathogens and biotechnologically relevant microbes

Nonulosonic acids (NulOs) are large nine-carbon sugars containing a highly acidic α-keto acid moiety. Beyond their distinctive chemistry, they often cap glycoconjugates and fulfil key biological functions. In humans, NulOs regulate protein half-life, and immunity^48^, while in microbes they can interfere with phage binding^49^ and are important virulence factors in pathogens, such as *Campylobacter jejuni* and *Helicobacter pylori*^*5,50*^. Their analysis is challenged by extensive chemical diversification and instability. However, understanding their diversity and biosynthetic pathways provides promising targets for the development of novel antimicrobials.

SugarBase identifies CMP-NulO derivatives based on diagnostic features: the CMP fragment, characteristic CO_2_ loss from the acid group, strong CMP dissociation due to the labile monophosphate ester, and a distinct nitrogen-rich composition^16^. To further explore the identified NulOs, another fraction of the metabolite extract was subjected to mild acid hydrolysis, labeled with 1,2-diamino-4,5-methylenedioxybenzene (DMB), and analyzed by DIA-MS as described by Kleikamp et al.^36^. This confirmed the presence of the α-keto acid, the carbon backbone length (e.g., C8, C9, or larger), the redox state (e.g., Neu versus Leg/Pse), and the nature of N- and O-linked modifications.

As these sugars are key modulators of cellular function, we investigated the CMP-NulO profiles of *Ca*. Accumulibacter phosphatis, *Magnetospirillum gryphiswaldense, Ca*. Kuenenia stuttgartiensis, and four *Campylobacter jejuni* strains. In *C. jejuni*, NulOs are O-linked to the FlaA and FlaB proteins and are essential for flagellar assembly and motility, with specific modifications influencing host cell invasion^2-5^. The strain NCTC 11168 is a well-characterized, highly invasive reference strain and a common cause of human gastroenteritis, whereas strain 81116 is less invasive and was first isolated during a waterborne outbreak in the UK^51-53^. NCTC 11168 produces both pseudaminic and legionaminic acid derivatives with a well-defined modification profile^2^, whereas strain 81116 lacks the legionaminic acid biosynthesis pathway^54^ and its NulO structures have not been characterized before. Across all these strains we identified over twenty distinct CMP-linked sugars (Figure 4a, SI Excel 6), of which all but two could be confidently assigned to ulosonic acid derivatives. *C. jejuni* 11168 showed the greatest CMP-NulO diversity, with at least eight distinct variants. Identified structures included NulOAcAc and NulOAcAm species, additional O-acetylation and methylation, 2,3-di-O-methylglyceroyl and methyl acetimidoyl modified (the latter has so far only been identified for Leg), consistent with previous reports for this strain^2^. Notably, the acetimidoylamino (Am; CH_3_C(=NH)-NH-) modification generated by the amidotransferase PtmG (Cj1324) predominated over the acetamido modification (Ac; CH_3_C(=O)–NH–) (SI Excel 6). This likely has functional implications, as protonation of the imine at physiological pH yields a net-neutral sugar, whereas the acetamido form is negatively charged, and contributes to a negative surface charge on proteins or cells. Interestingly, in addition to Pse/Leg acetimidoylamino modifications, we also observed peaks consistent with acetimidoylamino modified Neu. Notably, the nucleotide sugar pool predominantly contained the putative Neu5Am form, whereas after acid hydrolysis and DMB labeling, Neu5Ac was the predominant species. A similar shift from Am to Ac was also observed for Pse and Leg derivatives. This shift likely originates from reacetylation during acetic acid treatment and the amidotransferase may also preferably act on nucleotide sugars. Interestingly, we observed that acetimidoyl modifications may also affect other N-acetylated sugars, such as UDP-HexNAc, which has not been reported previously. However, their potential incorporation into glycoconjugates was not further investigated in this study.

**Figure 4.**
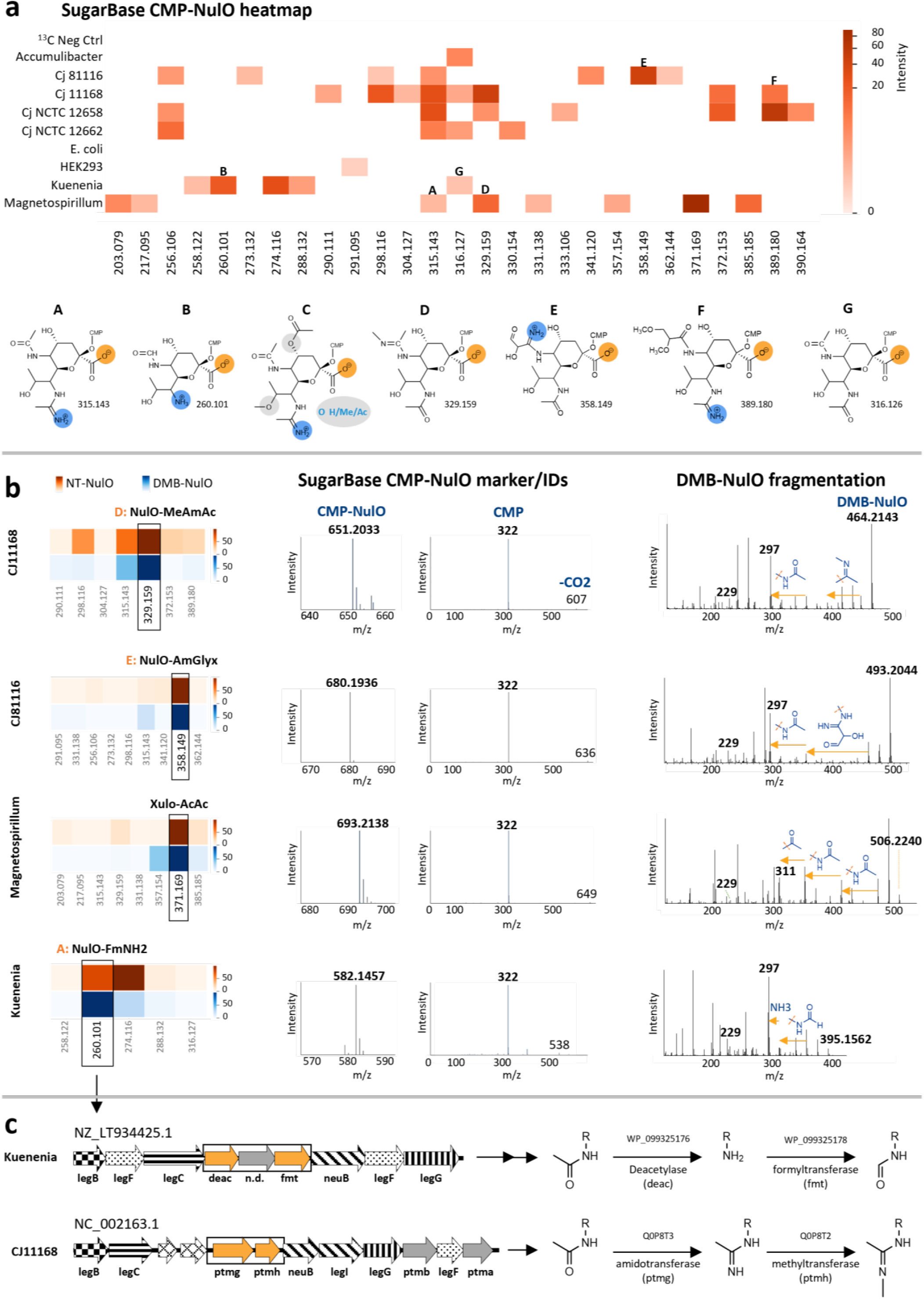
SugarBase CMP-NulO profile in pathogens and environmental microbes. **a)** The heatmap shows the SugarBase CMP-NulO profile across pathogenic *C. jejuni* and environmental microbes. The monoisotopic mass of identified NulOs are shown on the x-axis, and colors indicate normalized peak intensity. The schemes below present selected CMP-NulOs with their putative structures; no structure could be assigned for m/z 333. **b)** The graphs show SugarBase CMP-NulO identification and structural characterization using DMB labeling and fragmentation for the dominant NulO structures in *Campylobacter jejuni* 11168, *Campylobacter jejuni* 81116, *Magnetospirillum gryphiswaldense*, and *Ca*. Kuenenia stuttgartiensis. The middle panel shows SugarBase CMP-NulO pseudo-MS1 signals with characteristic CMP and CO_2_-loss fragments in MS2. The right panels show DMB labeling and fragmentation, confirming the ketoacid group, carbon-chain length and oxidation state, and modifications, with putative structures annotated. The left heatmaps compare the abundance of the CMP-NulOs and corresponding DMB-NulOs, with color indicating normalized intensity. **c)** The identified NulO type and modification in *Ca*. Kuenenia stuttgartiensis enabled identification of the corresponding biosynthetic gene cluster (genome NZ_LT934425.1). Notably, genes encoding a deacetylase and a formyltransferase, likely responsible for its unique modifications, are located between homologs of legB, legF, and legC, and putative neuB, legF, and legG genes. For comparison, the Leg biosynthetic pathway and methylation step from *C. jejuni* NCTC 11168 (genome NC_002163.1) are shown.

For *C. jejuni* 81116, the CMP-NulO modification profile has not been established previously. Using SugarBase, we observed a comparable structural diversity as for NCTC 11168, but here a single structure (m/z 680.1934, [M-H]^-^) accounted for approx. 85% of the total signal (Figure 4a & b; SI DOC). However, the inferred NulO moiety mass and sum formula (358.1488 Da, C1_4_H_22_N_4_O_7_) cannot be explained by simple acetylation or methylation of the canonical NulO backbone. The orthogonal DMB-NulO fragmentation revealed C9 Leg/Pse marker ions, an N-acetyl or acetimidoyl group, and an additional neutral loss of 102.04 Da, consistent with an N-modification of formula C_3_H_6_N_2_O_2_. Given the close genomic relatedness to NCTC 11168 and the observed mass and elemental composition, this modification likely represents a glyceroyl substituent on an acetimidoyl group, as present in NCTC 11168 (possibly lacking O-methylation and with one hydroxyl oxidized to an aldehyde, or a closely related variant). Further minor CMP-NulOs that were detected correspond to canonical NulOAcAm, O-acetylated, and dehydrated forms. Interestingly, two low-abundance species (<5% combined) were consistent with NulOAmNH_2_, featuring a free amine, including one variant lacking the N-acetyl group, likely arising from incomplete N-acetyltransferase activity or degradation.

NCTC12658 showed a CMP-NulO profile similar to 11168, including both smaller NulOAcAm and larger more hydrophobic 2,3-di-O-methylglyceroyl modified structures that were absent in NCTC12662. Strain 12662 showed a previously identified NulO monoisotopic mass of 330.154, likely corresponding to substitution of a hydrogen atom by an amino group (-NH_2_)^49^. Strain 12658 displayed an additional NulO mass of 333.106, which could not be annotated based on known structures, nor structurally elucidated from the available data. Both *C. jejuni* strains have also been investigated for their susceptibility to different phages^55^. Flagellotropic phages target the flagellum and require bacterial motility, where receptor recognition depends on flagellin O-glycosylation and thus may be sensitive to presence and type of NulOs^49,55-57^. Flagellotropic phages are of particular interest because by binding they attenuate motility and consequently virulence, making these phages especially promising antibacterial agents^57^.

The CMP-sugar profile of *Ca*. Accumulibacter was dominated by a single CMP-NulOAc2 species, whereas *M. gryphiswaldense* showed a surprisingly broad CMP-sugar profile (Figure 4a). In addition to low-abundant NulOAcAm and acetylated/methylated derivatives, we detected several large CMP-sugars ([M–H]^-^ m/z 679.1983, 693.2139, and 707.2296). These masses were initially consistent with additional acetylation and methylation, and efficient fragmentation with characteristic CO_2_ loss confirmed their identity as CMP-linked α-keto acids. However, fragmentation of the DMB-labeled compounds showed a different image. While the NulOAcAm and NulOAcAmOMe derivatives showed the expected C9 Pse/Leg marker fragment m/z 297.09, the larger species yielded the m/z 311.10 marker fragment, indicative of an extended carbon backbone (Figure 4a & b, SI DOC). This observation is consistent with earlier reports by Kleikamp et al., who described C10 marker fragments in *M. gryphiswaldense* ulosonic acids that could not be assigned to known structures^36^. Therefore, we established a fragmentation tree for the observed marker fragments (SI DOC), which showed sequential loss of water and ammonia, followed by one O-acetyl and two N-acetyl groups, ultimately yielding the m/z 311.10 carbon length marker fragment. Further isolation and MS^3^ fragmentation of m/z 311.10 revealed an initial CO/H_2_O loss, followed by stepwise cleavage of four carbon atoms to the m/z 229.06 core fragment (DMB-keto acid). In comparison, MS^3^ fragmentation of the DMB-LegAc_2_ m/z 297.09 marker and the DMB-Neu5Ac marker m/z 295.09 showed the same initial CO/H_2_O loss, followed by cleavage of only three carbon atoms to the same m/z 229.06 core fragment as observed in *M. gryphiswaldense*. This suggests the presence of an elongated carbon backbone and a previously unreported α-keto acid in *M. gryphiswaldense*. Moreover, O-methylation would be lost early during fragmentation, before extensive C–C bond cleavage. Interestingly, lower-abundance CMP-linked species with unusually low masses (m/z 203 and 217), consistent with HexNAc-related compounds, were also detected in *M. gryphiswaldense*, which furthermore were not detected in DMB-labelling experiments confirming the lack of an α-keto acid moiety.

Finally, we investigated *Ca*. Kuenenia stuttgartiensis, which was recently shown to express complex O-glycans containing NulOs with a free amino group. Like acetimidoyl modifications, free amines are protonated at physiological pH, yielding a net-neutral sugar despite the strong acidic α-keto group. Free amines have previously been reported only in certain cancer cells, where they influence cell surface charge and aggregation^58^. Consistent with these observations, the SugarBase workflow identified NulOs containing one free amine and a second amine that was formylated (rather than acetylated), and CMP-NulOs with +14.01 Da consistent with O-methylation (Figure 4a & b, SI DOC). The presence of these modifications in CMP-activated sugars show that deacetylation and formylation occur at the nucleotide sugar level and not after glycan biosynthesis. This prompted a closer examination of the *Ca*. Kuenenia stuttgartiensis genome to identify the gene cluster that enables the production of CMP-NulOs with an unsubstituted amine. We searched the genome for pseudaminic and legionaminic acid biosynthetic genes in proximity to enzymes such as deacetylases and methyl- or formyl-transferases. Indeed, we found a legionaminic acid-related biosynthetic gene cluster with putative legB, legF, and legC homologs, together with genes encoding a deacetylase and a formyltransferase, followed by putative neuB, legF, and legG genes (Figure 4c). This biosynthetic pathway intrinsically incorporates deacetylation and formylation steps. Although overall sequence homology is low, a similar genomic architecture is present in *C. jejuni* NCTC 11168, where ptmG and ptmH between legC and neuB mediate conversion of N-acetyl groups to acetimidoyl moieties and subsequent methylation (with the N-acetyltransferase legH located near the pseudaminic acid locus).

### SugarBase enables discrimination between strains that produce pseudaminic or legionaminic acid

Distinguishing nonulosonic acid (NulO) analogues with identical composition but different stereochemistry remains a major challenge. Among known NulOs, pseudaminic and legionaminic acids are the most common stereoisomers, while other variants are rarely reported^48^. Recently developed monoclonal antibodies specifically recognize Pse-containing glycoconjugates which allowed staining and glycoproteomic detection of Pse containing glycans^59^.

However, broader studies of this group of NulO molecules still require NMR and predictions based on genomic context. This is particularly challenging for hard-to-culture organisms and species with poorly annotated genomes.

Pseudaminic and legionaminic acids differ in stereochemistry at multiple chiral centers and are synthesized via distinct biosynthetic pathways^21,22,60^. Pseudaminic acid is derived from UDP-GlcNAc through sequential dehydration, epimerization, transamination, and acetylation steps. Nucleotidase activity then produces 2,4-diacetamido-2,4,6-trideoxy-L-altropyranose, which is condensed with phosphoenolpyruvate by Pse synthase to form Pse5Ac7Ac. This product is subsequently activated by the cytidylyltransferase PseF to generate CMP–Pse5Ac7Ac (Figure 5b, SI DOC). Legionaminic acid biosynthesis follows a similar pathway but begins with GDP-GlcNAc (Figure 5b), enabling metabolic partitioning and independent regulation of the two pathways^21,22,60^.

**Figure 5.**
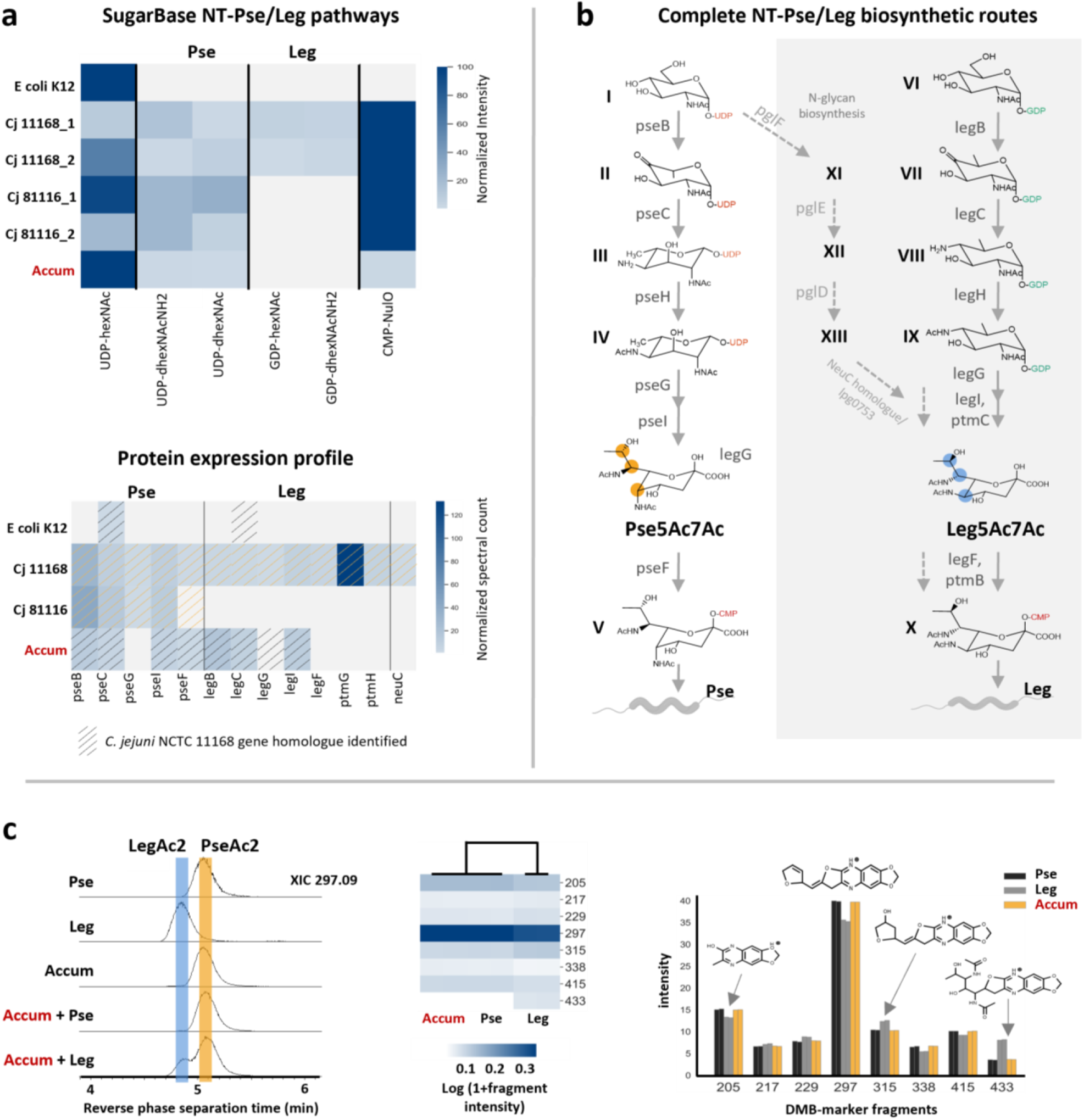
SugarBase discriminates between Pse and Leg producing strains. a) The upper heatmap shows SugarBase screening for key nucleotide sugar intermediates that distinguish Pse- and Leg-producing strains. For Pse, these include UDP-dHexNAcNH2 and UDP-dHexNAc; for Leg, GDP-HexNAc and GDP-dHexNAcNH2, shown together with the UDP-HexNAc precursor and CMP-NulO products. From top to bottom: *E. coli* K12, *C. jejuni* 11168 (duplicates 1–2), *C. jejuni* 81116 (duplicates 1–2), and *Ca*. Accumulibacter phosphatis. *E. coli* serves as a negative control, *C. jejuni* 81116 as a Pse-positive control, and *C. jejuni* 11168 as a Pse/Leg-positive control. SugarBase detects key intermediates in *Ca*. Accumulibacter phosphatis only for the Pse pathway. The lower heatmap shows proteomics data for the same strains, confirming expression of enzymes involved in Pse5Ac7Ac and Leg5Ac7Ac biosynthesis. Abundance is indicated by normalized spectral counts. The orange hatched pattern marks genes supported by genome annotation and literature^21,22,54^, while grey hatching indicates genes identified by sequence alignment to *C. jejuni* NCTC 11168 pathways. b) The schemes show biosynthesis pathways for Pse5Ac7Ac (from UDP-GlcNAc) and Leg5Ac7Ac (from GDP-GlcNAc). Intermediates are labeled I–XIII, covering key UDP- and GDP-linked sugar derivatives and final CMP-activated products. c) The left graph shows reverse-phase separation of DMB-Pse5Ac7Ac (Pse) and DMB-Leg5Ac7Ac (Leg), monitored by the C9 fragment (m/z 297.09). DMB-labeled *Ca*. Accumulibacter phosphatis (Accum) was analyzed with and without spiked standards. The Accum NulOAc2 co-elutes with the Pse marker. The heatmap (cosine similarity clustering) groups Accum fragments with the Pse standard. The bar plot shows diagnostic fragments that clearly distinguish Pse from Leg.

We used SugarBase to investigate the presence of Pse- and Leg-specific nucleotide sugar intermediates, which would indicate which of these compounds are produced. *C. jejuni* strain 11168 produces both Pse and Leg, whereas strain 81116 produces only Pse. Consistent with this, we observed nucleotide sugar intermediates for both Pse and Leg in *C. jejuni* NCTC 11168, while we only detected Pse-related intermediates in strain 81116 (Figure 5a, upper heatmap). We next examined *Ca*. Accumulibacter phosphatis, for which homology-based predictions and proteomics data provide only ambiguous evidence regarding the presence of Pse, Leg, or both pathways (Figure 5a, lower heatmap). This ambiguity largely reflects the low sequence homology of the associated enzymes across taxa. In contrast, nucleotide sugar intermediates show uniform chemical structures because modifications occur at the NulO product level. When investigating *Ca*. Accumulibacter with SugarBase, we detected only key intermediates of the Pse pathway (Figure 5a). To support this annotation, we performed acid hydrolysis and DMB labeling of *Ca*. Accumulibacter biomass, followed by separation and fragmentation analysis together with commercial Pse5Ac7Ac and Leg5Ac7Ac standards. Pse and Leg showed partial chromatographic separation and distinct fragment ion ratios, which confirmed that the DMB-NulO peak in *Ca*. Accumulibacter was Pse5Ac7Ac (Figure 5c, SI DOC). Here, separation and fragmentation of labelled NulOs were possible because only doubly acetylated forms were present and standards were available.

In summary, SugarBase enables differentiation between Pse- and Leg-producing strains based on key nucleotide sugar intermediates, without relying on genomic predictions or the need for large-scale fractionation for NMR studies. This provides a new strategy to profile microbial systems for Pse and Leg production.

## CONCLUSION

SugarBase enables fully untargeted mapping of microbial nucleotide sugar networks by screening crude cell extracts using porous graphitic carbon separation, narrow-window DIA, and a new algorithm with a large composition database comprising thousands of theoretical entries. This approach links nucleotide marker fragments to their corresponding nucleotide sugar-parent compounds. We also provide a comparative overview of monosaccharide classes and nucleotide activation preferences across a broad phylogenetic range of microbes, revealing extensive species-specific features and many high-confidence identifications that remain unannotated.

This study provides a quantitative map of the *C. jejuni* NulO repertoire and supports the identification of biosynthetic gene clusters, for example in *Ca*. Kuenenia stuttgartiensis, which produces NulOs with a free amine. Integration of orthogonal DMB-based fragmentation confirmed different nonulosonic acid classes, including N- and O-linked modifications that influence physicochemical properties relevant to microbial biology. Distinct NulO profiles in *Campylobacter* species may also affect susceptibility to flagellotropic phages. Notably, we also report the first evidence of previously undescribed CMP-activated higher-carbon ulosonic acids in *Magnetospirillum*. Finally, pathway-specific nucleotide sugar intermediates enabled discrimination between pseudaminic- and legionaminic-acid–producing microbes without requiring genomic information. This was supported by comparative chromatographic separation and fragmentation experiments using synthetic Pse and Leg standards.

Together, SugarBase provides a scalable approach to discover new sugar pathways and enzymes, accelerating glycobiology in non-model microbes and enabling access to chemically challenging nucleotide sugars for antimicrobial and vaccine development. The SugarBase pipeline is freely available as Python code and as a standalone desktop executable.

## Supporting information

SI DOC

## DATA AND CODE AVAILABILITY

The SugarBase pipeline is freely available as Python code and as a standalone desktop executable with a browser-based graphical user interface: https://sourceforge.net/projects/sugarbase-x/files. All acquired data are publicly accessible through the 4TU.ResearchData platform (https://data.4tu.nl project “SugarBase”). MATLAB functions are available upon request.

## ACKNOWLEDGEMENTS

The authors are grateful for valuable discussions with colleagues from the Department of Biotechnology and acknowledge funding from the Zero Emission Biotechnology program at TU Delft. We thank Dita Heikens for general support in the mass spec facility at the department of Biotechnology, and Rita Martins Costa for support in the cell culture laboratory at the Department of Bionanoscience. Dimitry Y. Sorokin for culturing and enabling the *M. gryphiswaldense* studies, Timmy Paez Watson for providing *Ca*. Accumulibacter phosphatis enrichment biomass, and all other colleagues who provided cell biomass for exploratory and screening experiments.

