## Supplementary material for "SugarBase: mapping glycomolecule precursors in microbes": SI DOC

#### A. SugarBase QuickStart and Overview

SugarBase is designed to process LC–MS/MS data (in .mzXML format) to identify nucleotide-activated sugars (NT-sugars) based on diagnostic nucleotide fragments and a predefined chemical composition space. The workflow combines pseudo-MS1 precursor information with MS2 fragment evidence, filters candidate masses using theoretical chemical compositions, and assigns a confidence score to each detected hit.

The pipeline supports both a discovery mode (screening a broad chemical space of possible nucleotide-sugars) and a targeted mode (searching only predefined compositions from an input list). The desktop executable allows to explore the results interactively, by providing the extracted ion chromatograms as well as the pseudo-MS1 and MS2 scans in which the potential NT-sugar was detected for each nucleotide-sugar hit. In addition, results can be exported as an Excel table.

Key analytical steps include MS1 preprocessing (deisotoping and filtering), MS2 fragment extraction, MS1–MS2 alignment, precursor matching against a theoretical chemical sugar space, detection of consistent signals across consecutive scans, and multi-parameter scoring to rank candidate nucleotide-sugar hits. SugarBase can be used to discover the nucleotide-sugar profile in microbial extracts without requiring any prior knowledge.

**QuickStart**

1. Download the desktop executable: https://sourceforge.net/projects/sugarbase-x/files/SugarBase_v08032026.zip/download
2. Open the desktop executable, see Figure S1.
3. Download the test data: https://sourceforge.net/projects/sugarbase-x/files/Test_data_SugarBase.zip/download
4. Select your input folder (Figure S1 → 1). A maximum of three .mzXML files is allowed per analysis.
5. Select your output folder (Figure S1 → 2).
6. Enter a sample name (Figure S1 → 4).
7. Select the analysis mode (Figure S1 → 5).
    a. 'Targeted' -> requires a '.xlsx' file which should include the chemical formulas of the sugars you want to target in the first column, e.g. C6H12O6 (Figure S1 → 3).
    b. 'Discovery' -> no Target.xlsx file is required.
8. To run the analysis with default parameters, click "Run Analysis". For a description of the input parameters, the reader is referred to the caption of Figure S1 or to the ReadMe file as provided on SourceForge (https://sourceforge.net/projects/sugarbase-x/files/).

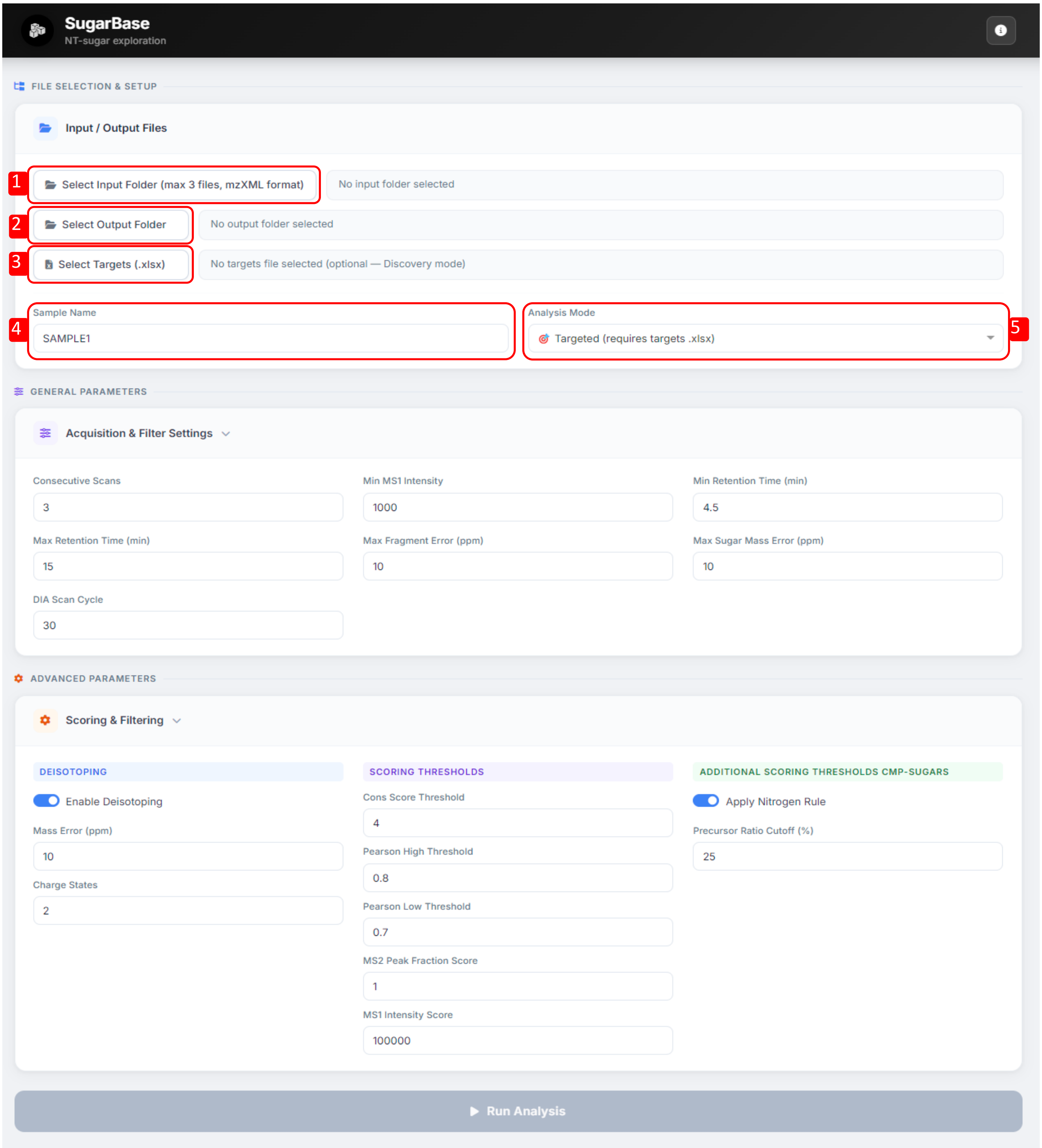

***Figure S1. Screenshot of the SugarBase desktop executable.*** For all parameters the default values are as shown in the Figure. Analysis mode: can be set to discovery (screen a broad chemical space of possible nucleotide sugars) or Targeted (only search for predefined compositions from an input list as defined in Select Targets ('xlsx'). GENERAL PARAMETERS. Consecutive Scans represents a filter parameter: hits representing potential nucleotide-sugars are only kept when the identified hit is present in at least X consecutive scans of the MS data. Min MS1 Intensity: a minimum intensity threshold for peaks identified in pseudo-MS1 spectra. Min Retention Time (min): a lower retention time cutoff (in minutes) for pseudo-MS1 data. Max Retention Time (min): an upper retention time cutoff (in minutes) for pseudo-MS1 data. Max Fragment Error (ppm): the mass accuracy (in ppm) for fragment ion searches. Max Sugar Mass Error (ppm): mass accuracy (in ppm) for MS1 data matching to the chemical sugar space. DIA Scan Cycle: Represents one PRM cycle. An 'DIA Scan Cycle' of 30 means that the same mass is targeted every $30^{th}$ scan, with consecutive scans at i, i+30, i+2*30, etc. ADVANCED PARAMETERS. Enable Deisotoping: whether to perform deisotoping of the pseudi-MS1 spectra. Mass Error (ppm): mass accuracy (in ppm) used for deisotoping. Charge States: charge state to consider when removing isotopic envelopes (e.g. 2 = remove up to doubly charged envelopes). Cons Score Threshold: a scoring parameter. A hit (potential nucleotide-sugar) scores

+10 when at least 'Cons Score Threshold' consecutive scans are present. Otherwise, the score is 0. Pearson High threshold: a scoring parameter. A hit scores +10 if the Pearson correlation of the precursors and fragments across three consecutive scans > 'Pearson High Threshold'. Pearson Low Threshold: a scoring parameter. A hit scores -10 if the Pearson correlation of the precursors and fragments across three consecutive scans < 'Pearson Low Threshold'. Note, the ‘Pearson Low Threshold’ should be >= 0.50, since a score of -50 is assigned when the Pearson correlation <0.50. Moreover, ‘Pearson High Threshold’ cannot be set smaller than ‘Pearson Low Threshold’. MS2 Peak Fraction Score: a scoring parameter. A hit scores +10 if the intensity ratio of the MS2 fragment compared to the "MS1" precursor > 'MS2 Peak Fraction Score' AND < 100. A score of -50 is assigned when the ratio > 110. MS1 Intensity Score: a scoring parameter. A hit scores +10 if the MS1 Intensity of the precursor > 'MS1 Intensity Score'. Otherwise, the score is 0. Apply Nitrogen Rule: Enables filtering of CMP-hit data using the nitrogen rule. According to the nitrogen rule, a valid CMP-NulO hit must have monosaccharides with both N and H atoms either even or odd in number; all others are excluded. Precursor Ratio Cutoff (%): Enables filtering of CMP-hit data based on the intensity ratio of the precursor identified in the MS2 scan compared to the fragment intensity in the MS2 scan. The red outlined numbers 1-5, correspond to references in the text section ‘QuickStart’.

#### B. SugarBase interactive dashboard interpreter

After running an analysis in SugarBase a tabular output will appear (Figure S2), presenting the identified nucleotide-sugar hits with various output parameters, such as total score, NT-Sugar and Sugar Mass. A maximum total score of 50 can be obtained, with 50 being very likely that the candidate is indeed a nucleotide-sugar. For a full list of output parameters, the reader is referred to the caption of Figure S2 or the ReadMe file provided on SourceForge (https://sourceforge.net/projects/sugarbase-x/files/). The tabular output can be exported as an Excel file by using the 'Export' function. By selecting one of the identified hits in the table, four graphs will appear, which allow for interactive exploration of the data. These graphs include: i) The extracted ion chromatograms of the precursor ion and the corresponding nucleotide fragment ion; ii) The three consecutive scans used to identify a putative nucleotide-sugar. Both the ion intensity of the precursor ion and of the fragment ion are displayed; iii) The MS2 spectrum of the identified hit; iv) The pseudo-MS1 spectrum of the identified hit. The graphs can be explored by using the cursor to zoom in and out.

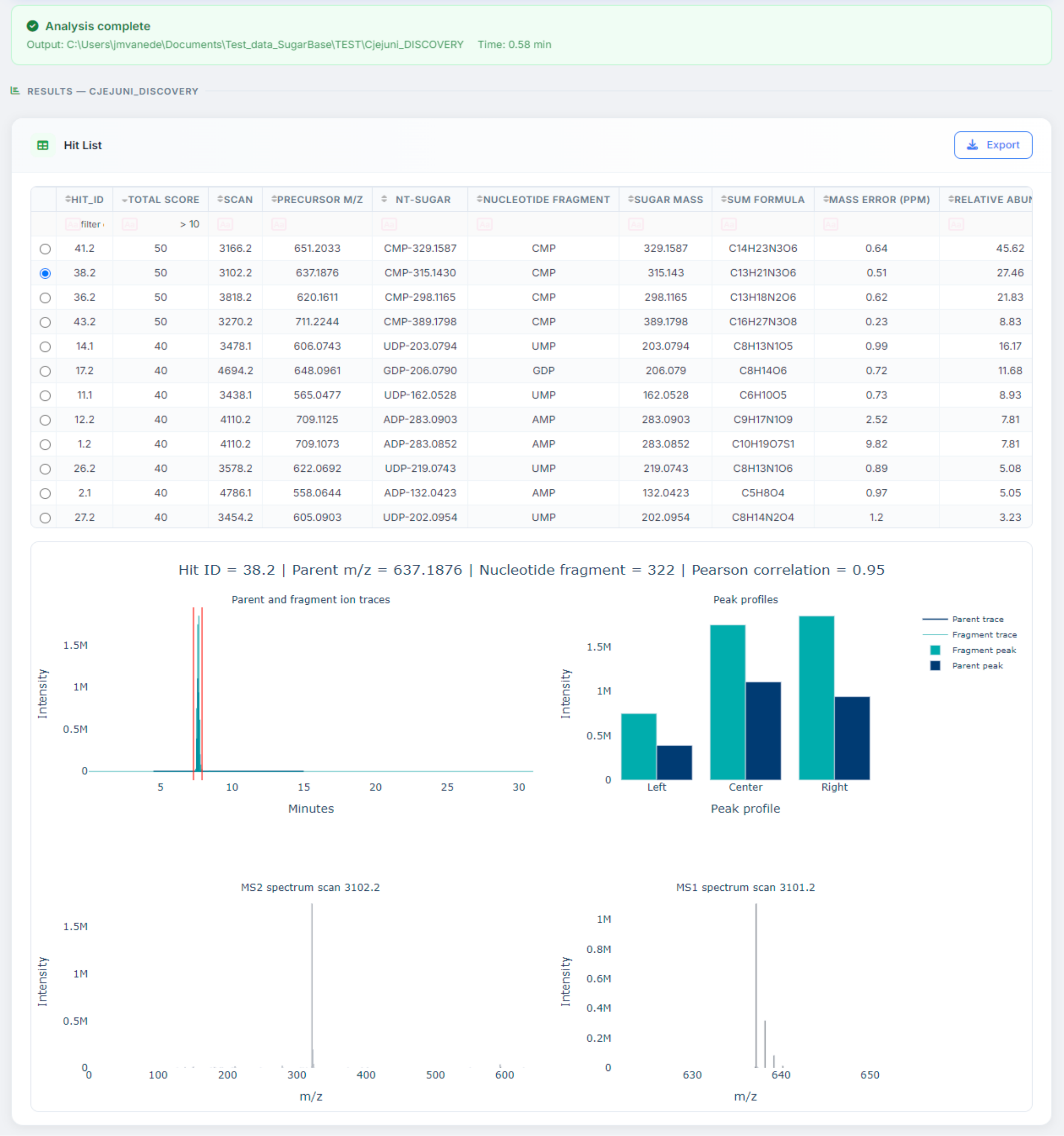

***Figure S2. Screenshot of the results of a C. jejuni 11168 sample analyzed by the SugarBase desktop executable in discovery mode using default parameters***. The Hit List provides a tabular overview of all the identified nucleotide-sugar candidates, which can be exported as an excel file using the 'Export' function. The table can be explored interactively by selecting one of the hits. This will result in four graphs appearing underneath the table output. The top left graph shows the extracted ion chromatograms of the precursor ion in the pseudo-MS1 scan and the nucleotide fragment ion in the MS2 scan. The top right graph, shows the three consecutive scans on which the hit is identified thereby displaying the peak intensity of the precursor ion of the pseudo MS1-scan (dark blue) and the fragment ion of the MS2 scan (turquoise). The bottom left graph shows the MS2 spectrum of the identified hit. The bottom right graph shows the pseudo-MS1 spectrum of the identified hit. The graphs can be explored by using the cursor to zoom in and out. The following parameters are displayed in the table: HIT_ID: A unique identifier assigned to each potential nucleotide-sugar hit. Total score: The overall score assigned to the identified hit. Scan: The scan number associated with the identified hit. Precursor m/z: The m/z value of the theoretical nucleotide-sugar, derived from the chemical composition database. NT-Sugar: The nucleotide combined with the monoisotopic mass of the corresponding monosaccharide. Nucleotide Fragment: The nucleotide fragment detected for the given hit. Sugar Mass: The monoisotopic mass of the identified potential monosaccharide. Sum Formula: The chemical sum formula of the potential monosaccharide. Mass Error (ppm): The difference (in parts per million) between the mass of the identified hit and the theoretical mass of the corresponding nucleotide-sugar from the chemical composition database. Relative abundance: The relative intensity (%) of the hit, normalized to the most intense hit detected within the same sample. MS1 Intensity: The MS1 signal intensity corresponding to the identified hit. RT (min): The chromatographic retention time of the hit, expressed in minutes. Isolation centre: The m/z center of the isolation window used during acquisition. Consecutive Score: Assigned +10 if the hit is detected in at least X consecutive scans; otherwise, the score is 0. Pearson Score: Assigned +10 if the Pearson correlation of the precursor and corresponding fragment intensities across three consecutive scans exceeds 'Pearson High Threshold'. A score of -10 is assigned if < 'Pearson Low Threshold' and a score of -50 when <0.50. Frag Score: Assigned +10 if the intensity ratio of the MS2 fragment relative to the "MS1" precursor is <100, a score of -10 is assigned when <'MS2 Peak Fraction Score' and a score of -50 when >110. Parent Score: Assigned +10 if the MS1 Intensity of the identified hit exceeds 'MS1 Intensity Score'; otherwise, the score is 0. CO2 Score: Assigned +10 if a CO2 loss peak is present in the corresponding "MS2" spectrum of the identified hit; otherwise, the score is 0. MS1/MS2 Correlation: the Pearson correlation of the precursors and fragments across three consecutive scans. % Fragmentation: The MS2 fragment intensity/MS1 precursor intensity (or MS2 precursor intensity for CMP-sugars)*100%

#### C. Theoretical nucleotide sugar chemical composition space.

***Table S1***. Filter criteria for the theoretical nucleotide-sugar chemical composition space.

| | Remove compositions with: | |
|---|---|---|
| Mass | A mass | < 100 |
| | A mass | > 400 |
| Mass defect | A mass defect | < 0.01 |
| | A mass defect | > 0.19 |
| | A mass between 100-200 and a mass defect | > 0.09 |
| | A mass between 100-250 and a mass defect | > 0.13 |
| Double Bond Equivalent (DBE) | A DBE | < 1.0 |
| | A DBE | > 6.0 |
| | A mass between 100-150 and a DBE | > 3.0 |
| C/H Ratio | A C/H ratio | < 0.40 |
| | A C/H ratio | > 1.25 |
| C/(O+N) Ratio | A C/(O+N) ratio | < 0.90 |
| | A C/(O+N) ratio | > 2.10 |
| Nr. of N- and S atoms | A mass between 100-200 and nr. of N-atoms | > 1 |
| | A mass between 100-275 and nr. of N-atoms | > 3 |
| | Nr. of S- and N-atoms | > 0 |

#### D. SugarBase scoring and filtering criteria.

***Table S2***. Scoring criteria for the identified NT-sugar hits.

| Scoring criteria | Threshold | Score |
|---|---|---|
| Number of consecutive scans in which the hit is present | >= 4 | +10 |
| Pearson correlation (based on three scans) | > 0.80 | +10 |
| | < 0.70 | -10 |
| | < 0.50 | -50 |
| For non-CMP-sugar hits: Percentage of the precursor intensity in the “MS1” scan compared to the fragment intensity in the “MS2” scan. | < 100 | +10 |
| | < 1 | -10 |
| | > 110 | -50 |
| For CMP-sugar hits: Percentage of the precursor intensity in the “MS2” scan compared to its intensity in the “MS1” scan | <= 25 | +10 |
| | > 25 | -50 |
| Precursor intensity in the “MS1” scan | > 100000 | +10 |
| Presence of a CO2 loss peak in the “MS2” scan | Pass | +10 |
| For CMP-sugar hits: N-rule: contains both an even or both an odd number of N- and H-atoms | Fail | -50 |

#### E. Retention time variability of the porous graphitic carbon column.

Retention time accuracy of the porous graphitic carbon separation was monitored by injecting a $^{13}$C-labelled *Penicillium chrysogenum* reference metabolite extract prior to every sample batch (each batch containing a maximum of six samples).

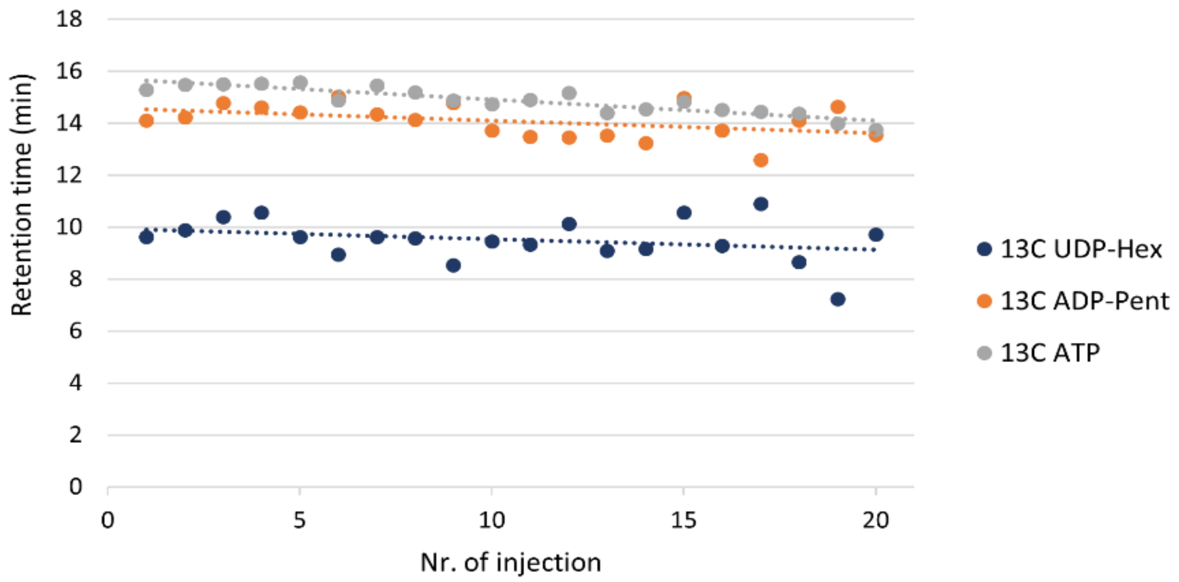

***Figure S3.*** **Retention time variability of the porous graphite separation.** The retention time of $^{13}$C UDP-hexose, $^{13}$C ADP-Pentose and $^{13}$C ATP is plotted against the date of injection date.

### F. Nucleotide sugar profile across analyzed taxonomic space of microbes.

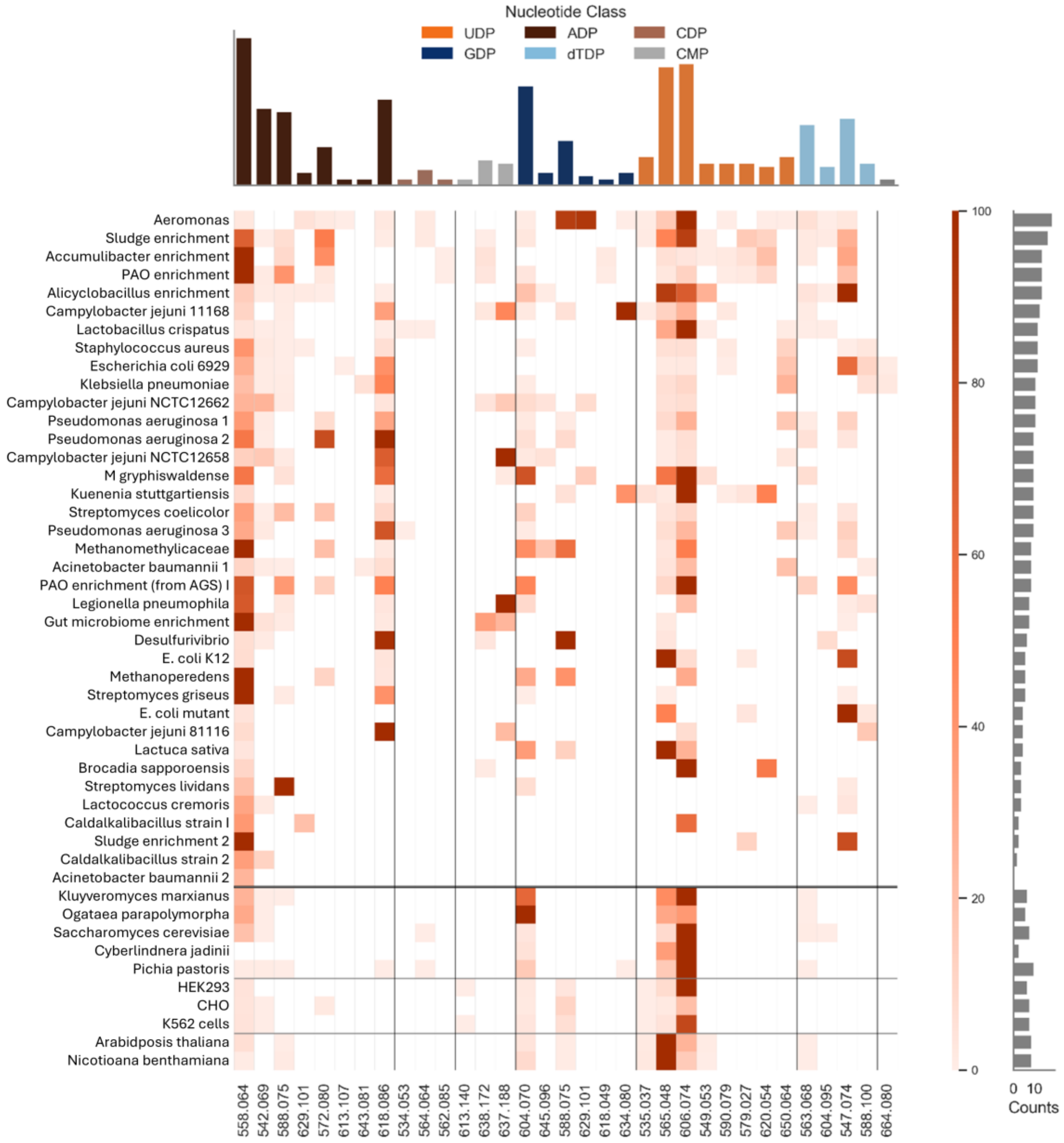

***Figure S4-1.*** Heatmap showing identified nucleotide-sugar candidates across diverse microorganisms using the established "SugarBase" pipeline in targeted mode. The pipeline was applied without an abundance threshold, but with the default filtering and scoring parameters. Nucleotide sugar candidates identified in >1 sample are displayed. The m/z values of the identified nucleotide sugar candidates ($[M–H]^–$) are displayed on the x-axis, and the color scale represents normalized peak intensities. The bar graph above the heatmap indicates the number of analyzed microorganisms in which each nucleotide sugar candidate was identified, while the bar graph on the right shows the number of nucleotide sugar candidates detected per microorganism.

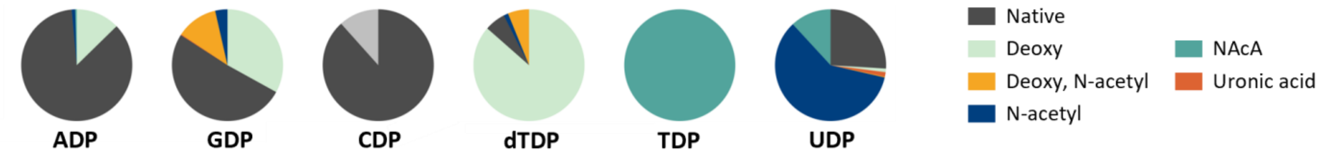

***Figure S4-2**.* The distribution of common sugar modifications across the different nucleotides-sugars identified over a broad phylogenetic spectrum of microbes. A matrix of 174 nucleotide sugars, linking common pentose, hexose, heptose, and nonulosonic acid sugars and their modifications to the seven activating nucleotides UDP, ADP, GDP, CDP, CMP, dTDP, and TDP, was queried using the SugarBase pipeline. The analyzed organisms are illustrated in Figure S4.

#### G. DMB-labelling reaction of α-keto acid group in ulosonic acids.

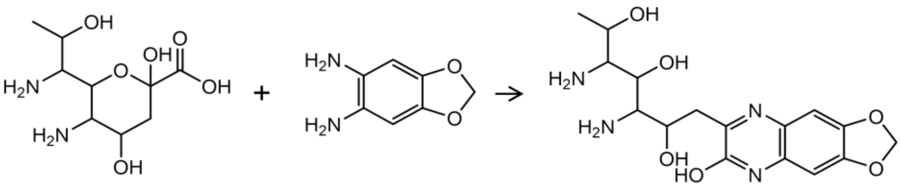

NulOAc2 DMB DMB-NulOAc2

***Figure S5.*** Chemical DMB-labelling reaction of α-keto acids.

#### H. Fragmentation spectra of hydrolyzed and DMB-labelled nucleotide-sugar extracts.

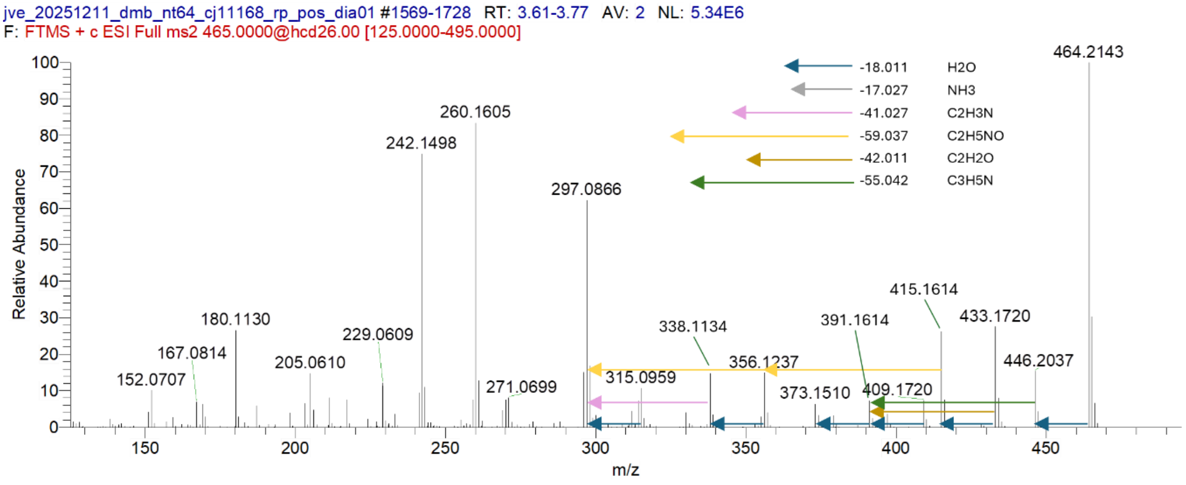

***Figure S6.*** Annotated fragmentation spectrum of m/z 464.2143 of a DMB-labelled nucleotide-extract from *Campylobacter jejuni* NCTC 11168.

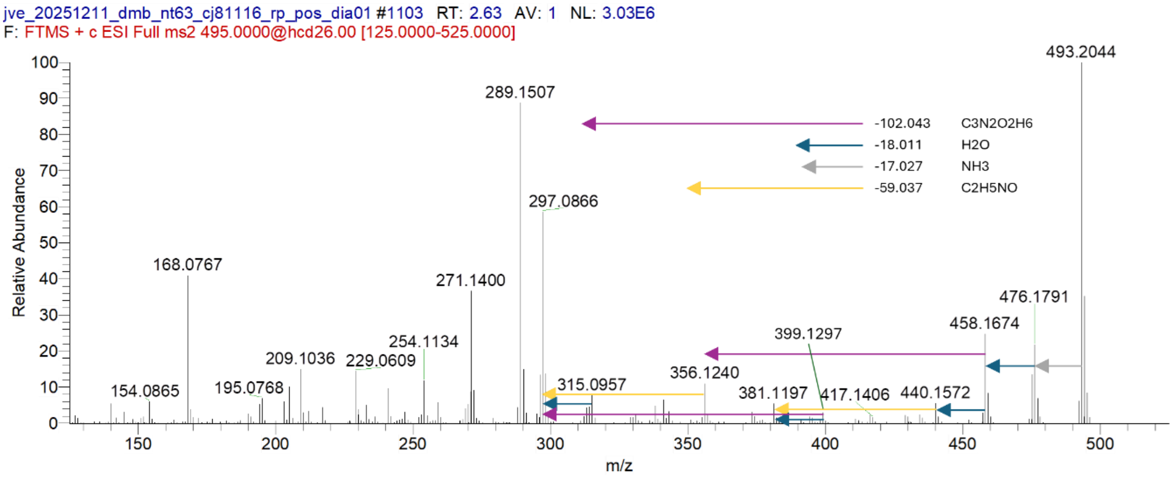

***Figure S7**.* Annotated fragmentation spectrum of m/z 493.2044 a DMB-labelled nucleotide-extract from *Campylobacter jejuni* NCTC 81116.

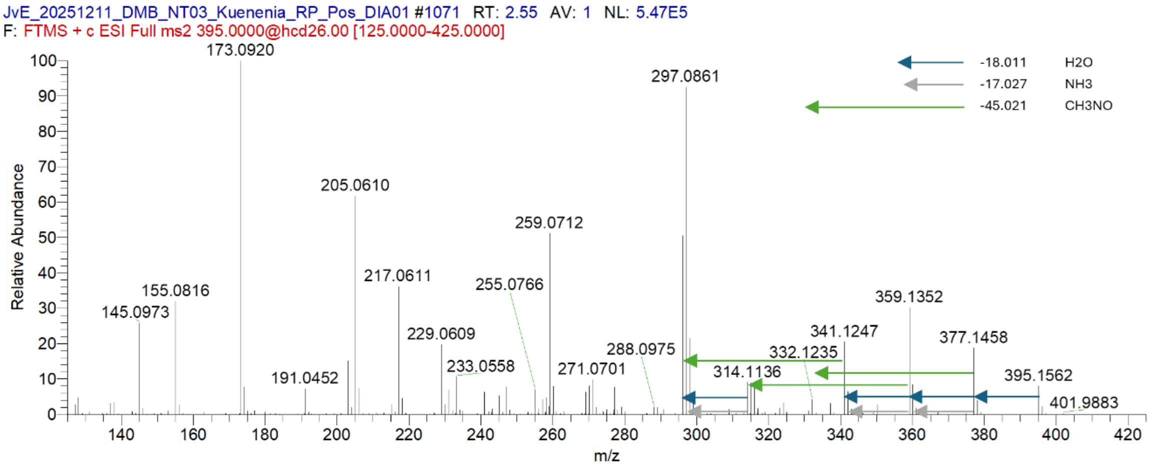

***Figure S8***. Annotated fragmentation spectrum of m/z 395.1562 of a DMB-labelled nucleotide-extract from *Candidatus* Kuenenia stuttgartiensis.

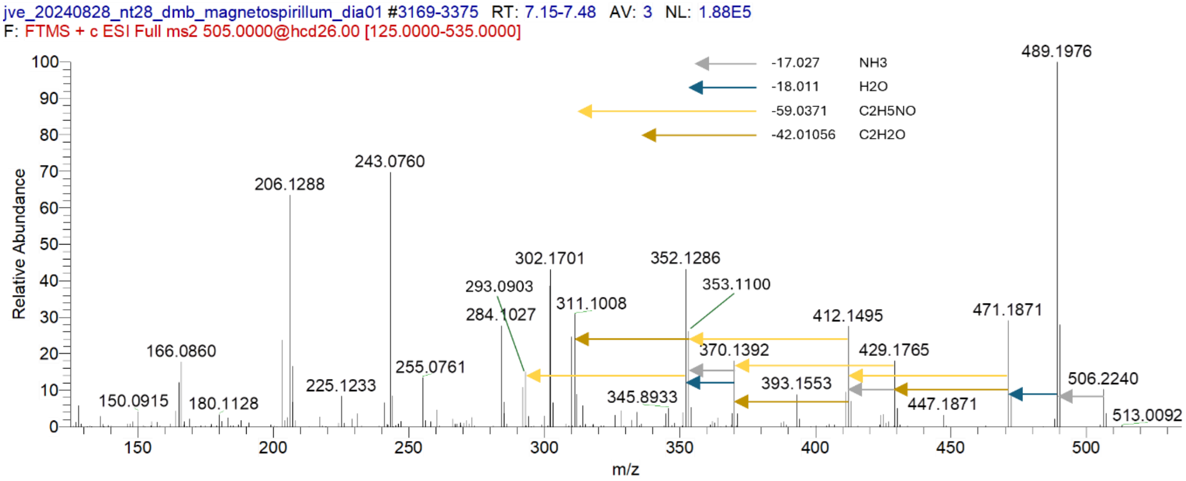

***Figure S9***. Annotated fragmentation spectrum of m/z 506.2240 of a DMB-labelled nucleotide-extract from *Magnetospirillum gryphiswaldense*.

#### I. MSn fragmentation tree of DMB-labelled higher carbon (C10) and C9 ulosonic acids

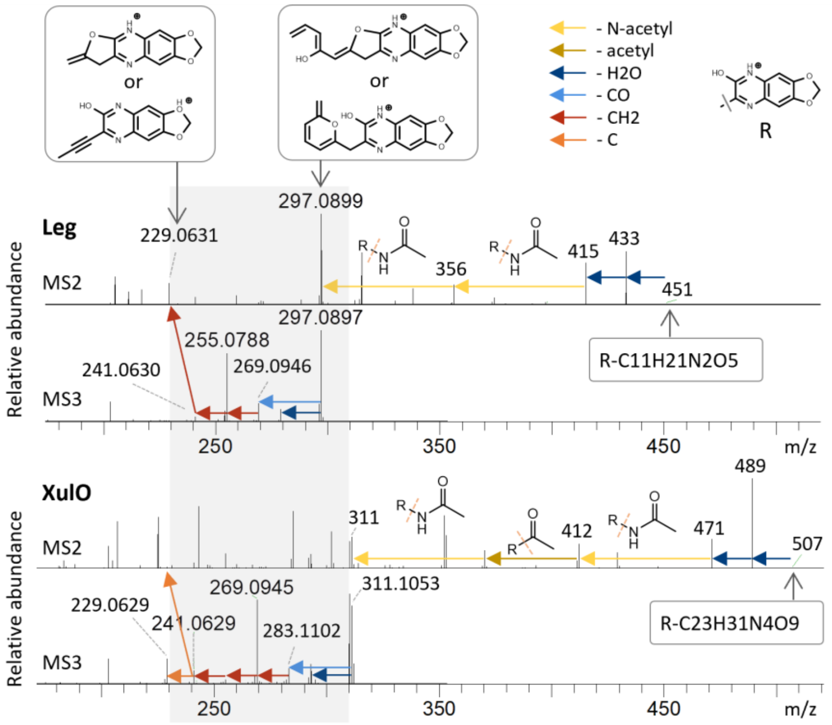

***Figure S10*. Fragmentation tree of the CMP-activated (C10) ulosonic acid in *M. gryphiswaldense*.** The graphs show the fragmentation trees of DMB-Leg5Ac7Ac (Leg) and the DMB-labeled higher-carbon ulosonic acid from *M. gryphiswaldense*, termed “XulO” (deculosonic acid). For Leg, fragmentation of the precursor ion (m/z 451) produced the characteristic C9 marker fragment at m/z 297.089, which was further isolated and fragmented to yield a fragmentation tree terminating in the conserved core fragment at m/z 229.06. Similarly, fragmentation of the XulO precursor ion (m/z 507) generated a C10 marker fragment at m/z 311.105, which upon isolation and fragmentation showed the same fragmentation tree, except fragmentation of one additional $CH_2$ unit before converging on the same m/z 229.06 core fragment. Fragment losses are annotated by coloured arrows.

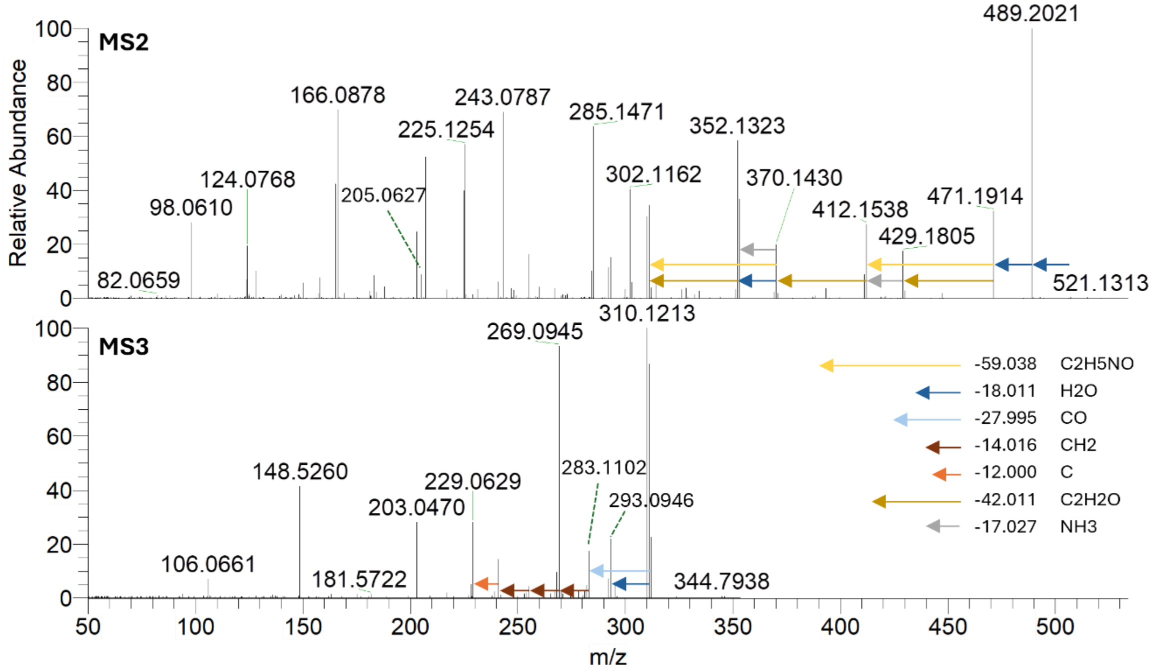

***Figure S11*.** Annotated fragmentation spectrum DMB labelled *Magnetospirillum gryphiswaldense.* Top: full MS2 spectrum of m/z 507.2292 at an HCD of 25. Bottom: full MS3 spectrum of m/z 311.1053 at an HCD of 32.

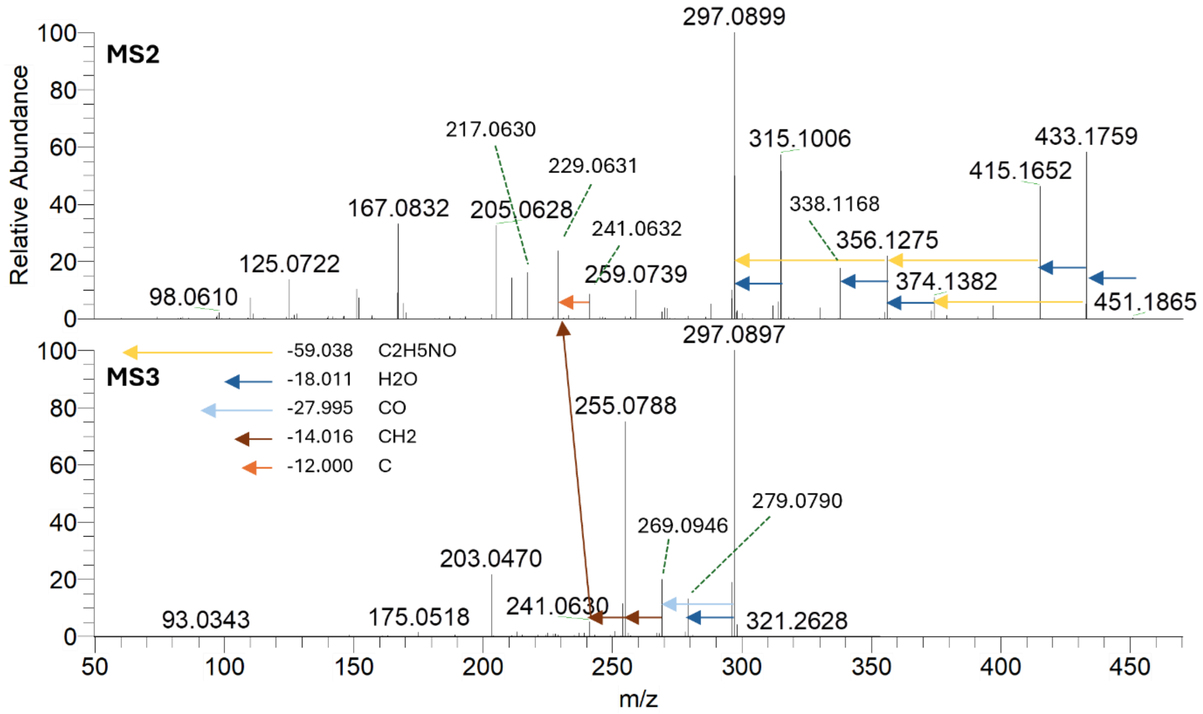

***Figure S12***. Annotated fragmentation spectrum DMB labelled Leg5Ac7Ac. Top: full MS2 spectrum of m/z 451.1650 at an HCD of 25. Bottom: full MS3 spectrum of m/z 297.0895 at an HCD of 32.

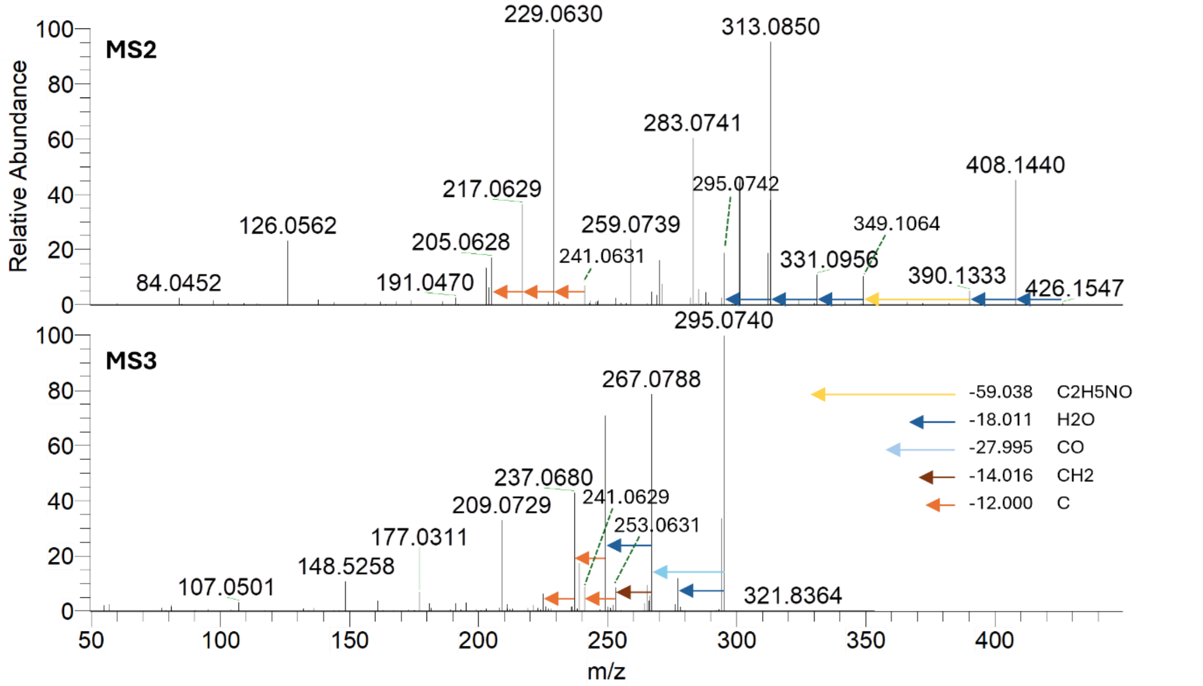

***Figure S13.*** Annotated fragmentation spectrum DMB labelled Neu5Ac. Top: full MS2 spectrum of m/z 426.1398 at an HCD of 25. Bottom: full MS3 spectrum of m/z 295.0739 at an HCD of 32.

#### J. Fragmentation of DMB-labelled *Ca.* Accum phosphatis biomass.

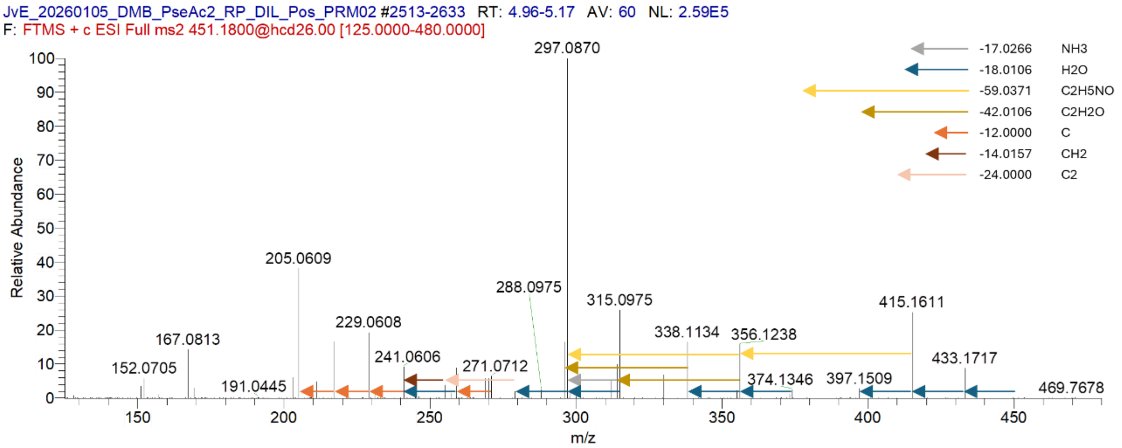

***Figure S14.*** Annotated fragmentation spectrum of m/z 451.180 of DMB-Pse5Ac7Ac.

#### K. DMB-Pse5Ac7Ac and DMB-Leg5Ac7Ac fragmentation profiling

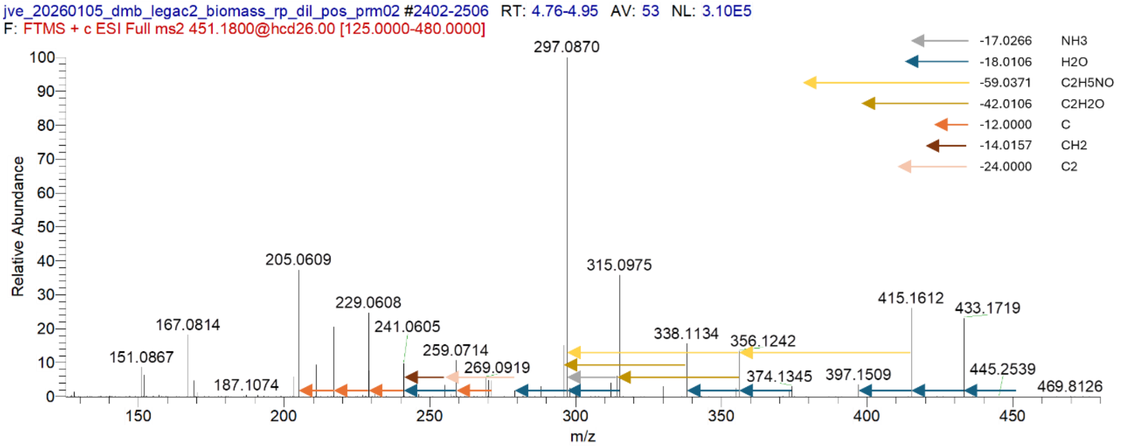

***Figure S15****.* Annotated fragmentation spectrum of m/z 451.180 of DMB-Leg5Ac7Ac.

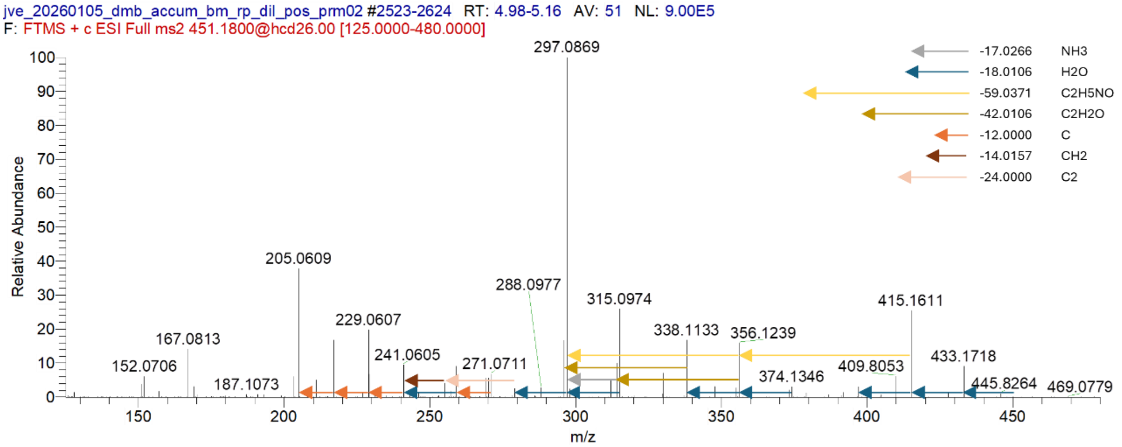

***Figure S16****.* Annotated fragmentation spectrum of m/z 451.180 from DMB-labelled *Ca.* Accumulibacter phosphatis biomass.

***Table S3***. Integrated area of DMB-Pse5Ac7Ac and DMB-Leg5Ac7Ac fragments with a 10 ppm error marge.

| | PSE1 | PSE2 | LEG1 | LEG2 | Accum1 | Accum2 |
|---|---|---|---|---|---|---|
| 205.1 | 103600 | 106660 | 120670 | 120119 | 351222 | 342889 |
| 217.1 | 45539 | 46630 | 64196 | 66200 | 157291 | 150912 |
| 229.1 | 53729 | 53653 | 80018 | 79774 | 185671 | 179444 |
| 297.1 | 275079 | 279266 | 322017 | 320514 | 930397 | 904737 |
| 315.1 | 71617 | 72628 | 111823 | 114555 | 240041 | 234355 |
| 338.1 | 44584 | 46575 | 49206 | 49927 | 157403 | 152379 |
| 415.2 | 69643 | 71003 | 83184 | 83620 | 235644 | 231251 |
| 433.2 | 24595 | 24548 | 73030 | 74634 | 85645 | 82553 |

#### L. Biosynthesis pathways of Pse5Ac7Ac and Leg5Ac7Ac.

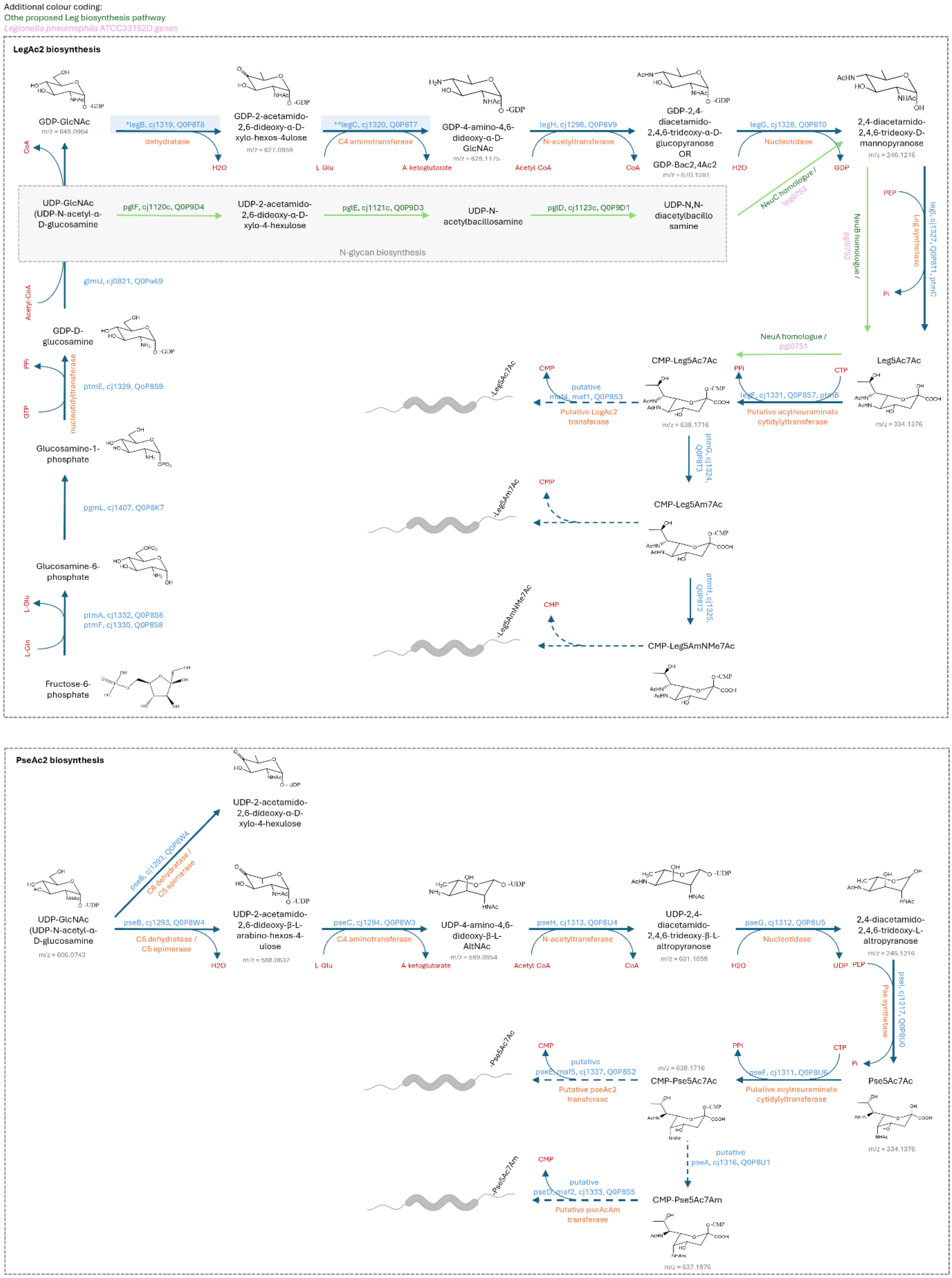

***Figure S17.*** Legionaminic acid and Pseudaminic acid biosynthesis pathways, including putative transferases[1-6].
